# Chromosome-scale *de novo* diploid assembly of the apple cultivar ‘Gala Galaxy’

**DOI:** 10.1101/2020.04.25.058891

**Authors:** Giovanni A.L. Broggini, Ina Schlathölter, Giancarlo Russo, Dario Copetti, Steven A. Yates, Bruno Studer, Andrea Patocchi

## Abstract

Apple (*Malus* × *domestica*) is one of the most important fruit crops in terms of worldwide production. Due to its self-incompatibility system and the long juvenile period, breeding of new apple cultivars combining traits desired by growers (e.g. yield, pest and disease resistance) and consumers (e.g. fruit size, color, and flavor) is a long and complex process. Genomics-assisted breeding strategies can facilitate the selection of germplasm leading to new cultivars. While the most complete apple genome assemblies available to date are from anther-derived homozygous lines, *de novo* assembly of apple genomes encompassing the natural heterozygosity remains challenging. Using long- and short-read sequencing technologies in combination with optical mapping, we *de novo* assembled a diploid and heterozygous genome of the apple cultivar ‘Gala Galaxy’. This approach resulted in 154 hybrid scaffolds (N50 = 34.3 Mb) spanning 999.9 Mb and in 414.7 Mb of unscaffolded sequences. Anchoring 31 scaffolds with a genetic map was sufficient to represent an entire haploid genome of 17 pseudomolecules (719.4 Mb). The remaining sequences were assembled in a second set of 17 pseudomolecules, which spanned 601 Mb, leaving 80.6 Mb of unplaced sequences. A total of 41,264 genes were annotated using 74,900 transcripts derived from RNA sequencing of pooled leaf tissue samples. This study provides a high-quality diploid reference genome sequence encompassing the natural heterozygosity of the widely popular cultivar ‘Gala Galaxy’. The DNA sequence resources and the assembly described here will serve as a solid foundation for fundamental and applied apple breeding research.

## Introduction

Apple (*Malus* × *domestica*) is the third most important fruit crop worldwide, with an annual production reaching 83 million metric tons in 2017 (FAOSTAT 2004). Most of the commercially successful cultivars are susceptible to a number of pests and diseases (Turechek 2004), and the production of high-quality fruits requires many applications of plant protection products. Given the increasing public demand to reduce fungicide input, disease resistant cultivars are essential for future sustainable apple production. Genotypic information has the potential to speed up the development of resistant cultivars of superior quality (Baumgartner, et al. 2016; Laurens, et al. 2018; Peace, et al. 2019). However, the number of available molecular markers linked to fruit quality traits is low. Several projects using genome-wide association studies or establishing genomic selection for fruit quality traits in apple were initiated (Kumar, et al. 2012; Muranty, et al. 2015; Roth, et al. 2019), often relying on SNP chip-based genotypic data (Bianco, et al. 2016; Bianco, et al. 2014). Once the association of genetic markers to a trait is observed, a complete reference genome sequence is a requisite for linking these markers to the underlying genomic features. With decreasing per base sequencing costs, sequencing-based genotyping (e.g. genotyping-by-sequencing or skim sequencing) requiring a reference genome for sequence alignment may partly supplant SNP chip-based genotyping. To meet the challenges described above, several projects already sequenced apple genomes (Daccord, et al. 2017; Li, et al. 2016; Velasco, et al. 2010; Zhang, et al. 2019). The *de novo* assembly of heterozygous genomes is recognized as a challenging task, as they generally result more fragmented than homozygous genomes (Pryszcz and Gabaldon 2016). So far, Velasco (2010) and Li (2016) assembled the genome of the heterozygous genotype ‘Golden Delicious’, with both assemblies resulting fragmented (N50 = 16 kb and 111 kb, respectively). Daccord *et al*. (2017) and Zhang *et al*. (2019) overcame the assembly challenges by sequencing the genome of the homozygous anther-derived lines GDDH13 and HFTH1 obtaining more contiguous assemblies (N50= 5.5 Mb and 7.0 Mb, respectively).

A complete heterozygous genome assembly allows investigating haplotype divergence within a single cultivar, but also to study the variation in the genome of sport mutants from a specific apple cultivar. Sport mutants are variations identified as single branches of the original cultivar producing fruits with improved characteristics (Foster and Aranzana 2018). For instance for the popular cultivar ‘Gala’, more than 30 sport mutants have been commercialised (Dickinson and White 1986; Iglesias, et al. 2008) and represent one of the most successful cultivar group worldwide. The exact reason for the emergence of sport mutants is unclear but likely due to imperfect repair following errors in DNA replication, active transposable elements (Foster and Aranzana 2018; Lee, et al. 2016) and/or epimutations (inheritable changes of DNA methylation, histone acetylation or chromatin remodeling) that influence genes transcription (El-Sharkawy, et al. 2015). Therefore, the availability of a complete heterozygous (diploid) genome assembly, besides supporting conventional breeding research, can help understanding the events underlying the generation of novel sport mutants and, more generally, the accumulation of mutations in this vegetatively propagated crop.

## Material and Methods

### Assembly strategy

Genomic DNA was isolated from field-grown apple leaves of ‘Gala Galaxy’ and processed for library construction for long- and short-reads sequencing as described in the supplementary methods. Additional long-reads libraries were also generated from four genotypes of the ‘Gala’ sport mutants group; ‘Gala’ original, ‘Gala Royal’, ‘Gala Schnico Red’ and the cisgenic apple line C44.4.146 (Kost, et al. 2015). The long-reads libraries from ‘Gala Galaxy’ and the additional genotypes were sequenced using PacBio RSII and Sequel instruments, respectively. The complete PacBio sequence dataset was *de novo* assembled with FALCON Unzip, v.0.4.0 (Chin, et al. 2016). Contigs consisting in more than 50% organellar (chloroplast or mitochondrial) genome sequences were identified by BLAST search (Altschul, et al. 1990) and removed, together with contigs smaller than 1 kb. Short-reads libraries (‘Gala Galaxy’) were sequenced on an Illumina Hiseq4000 using the paired-end 150 bp module. These were used for polishing of the assembly and for genome size estimation as described in the supplementary methods.

Contiguity of the assembly was then increased combining two optical maps to generate dual enzyme hybrid scaffolds. DNA extraction and labelling, and assembly strategy of the optical maps are described in the supplementary methods. The scaffolds were then oriented and ordered by ALLMAPS (Tang, et al. 2015) to achieve a chromosome-scale assembly, as described in the supplementary methods. Several ALLMAPS runs integrated the two evidence sets producing chromosome-scale pseudomolecules for both sets of homologous chromosomes. The diploid assembly was investigated for completeness using the Benchmarking Universal Single-Copy Orthologs (Simao, et al. 2015) pipeline v3.0 with 1,440 conserved plant genes (embryophyta_odb9) available in the discovery environment of www.cyverse.org. To evaluate the correctness of the genome assembly, diagnostic genome features were visualized in RStudio (Version 1.2.5001) using the karyoploteR package (Gel and Serra 2017) as described in the supplementary methods.

### Gene Annotation

For evidence-based gene annotation, RNA was extracted separately from 15 pools of three leaves collected from 1-year old field-grown ‘Gala Galaxy’ trees. Leaves were frozen in liquid nitrogen and ground to fine powder. RNA extraction protocol and libraries preparation are described in the supplementary methods. Libraries were sequenced on an Illumina HiSeq4000 instrument, generating paired-end reads (length = 150bp). Reads were checked for quality using FastQC (http://www.bioinformatics.babraham.ac.uk/projects/fastqc/) and trimmed using fastp (Chen, et al. 2018) retaining only sequences with Q>30. Libraries were mapped against the reference genome using bowtie2 v2.2.3 (Langmead 2010) and transcripts were identified with cufflinks v2.1 and tophat 2.0.13 (Trapnell, et al. 2012). Transcripts were functionally annotated as described in Knorst et al. (2019). Gene annotations were used to investigate collinearity between haploid assemblies by generating synthenic dotplots with Synmap2 (Haug-Baltzell, et al. 2017).

### K-mer Analysis Toolkit

The use of a haploid reference for sequencing read mapping may result problematic if one or both haplotypes from a heterozygous genotypes show high divergence to the corresponding haplotype in the reference. Sequence reads from diverging haplotypes would not be mapped to the reference and result in information loss. The k-mer analysis toolkit (Mapleson, et al. 2016) was used to assess the presence and count of Illumina sequencing data (in k-mers) of ‘Gala Galaxy’ (heterozygous) in different assemblies. For this analysis, only forward (R1) Illumina reads (‘Gala Galaxy’) were used in the KAT analysis with the GDDH13 assembly (Daccord, et al. 2017) as well as the diploid assembly reported here as reference. In addition, we included in this analysis a graph-based phased genome generated as described in the supplementary methods with Whatshap (Patterson, et al. 2015) using the primary haploid assembly MDGGph_v1.0 as reference.

## Results and Discussion

### Sequencing

For the genotype ‘Gala Galaxy’, PacBio RSII generated a total of 32 Gb long-reads sequence data using 29 SMRT cells, while Illumina HiSeq4000 instrument generated 694 million short-reads (104 Gb, submitted to SRA, accession XXXXXX). Using 19-mers, this latter data allowed estimating a genome size of 710.5 Mb (data not shown). To increase coverage of long-reads sequences, PacBio Sequel generated additional long-reads sequence data for other four genotypes of the ‘Gala’ group; ‘Gala’ original (5 SMRT cells, 9.4 Gb), ‘Gala Royal’ (8 SMRT cells, 8.8 Gb), ‘Gala Schniga® SchniCo red’ (2 SMRT cells, 8.3 Gb) and C44.4.146 (3 SMRT cells, 11.6 Gb). Combining all generated long-reads data, FALCON unzip produced 2,061 primary contigs and 5,663 haplotigs (N50 = 0.59 Mb, total length = 1,308.97 Mb, Table 1). A total of 201 organellar contigs (total of 8.57 Mb) and 31 contigs smaller than 1 kb were removed from the assembly.

**Table 1:** Statistics of the different assemblies generated in this work.

|  | **Falcon Unzip** | **Two-enzymes**<br>**hybrid assembly** | **MDGGph_v1.0**<br>**(primary haploid**<br>**assembly)** | **MDGGsh_v1.0**<br>**(secondary**<br>**haploid**<br>**assembly)** | **MDGGdi_v1.0**<br>**(diploid**<br>**assembly)** |
| --- | --- | --- | --- | --- | --- |
| **Total number of**<br>**contigs/scaffolds** | 2,061 primary<br>contigs and<br>5,663 haplotigs | 154 hybrid<br>scaffolds and<br>6336 unscaffolded<br>contigs | 31 hybrid scaffolds<br>in 17<br>pseudomolecules | 3,662 hybrid<br>scaffolds and<br>contigs in 17<br>pseudomolecules<br>and 2583<br>unanchored<br>contigs | 34<br>pseudomolecules<br>from<br>MDGGph_v1.0<br>and<br>MDGGsh_v1.0<br>and 2583<br>unanchored<br>contigs |
| **Total length** | 1,308.969 Mb | 1,414.649 Mb<br>(999.96 Mb in<br>scaffolds + 414.68<br>Mb unscaffolded | 719.450 Mb | 601.426 Mb +<br>80.621 Mb of<br>unanchored<br>sequences | 1,401.497 Mb |
| **N50 length** | 0.585 Mb | 13.704 Mb<br>(34.286 Mb for<br>hybrid scaffolds<br>only) |  |  |  |
| **Ns** |  |  | 29,755,402 bp | 70,500,027 bp | 100,255,429 bp |

### Scaffolding

Two optical maps were used for scaffolding of the assembled contigs. The alignment of 316,748 (out of 2,290,666) NLRS-labelled molecules produced 1,662 optical contigs (N50 = 0.65 Mb, total length = 948.63 Mb, coverage = 20×). The alignment of 329,921 (out of 2,117,673) DLE-1 labelled input DNA molecules produced 296 optical contigs (N50 = 15.53 Mb, total length = 1,806.45 Mb, coverage = 29×). Dual enzyme hybrid scaffolding was performed using Bionano Access v1.2.1 and Bionano Solve v3.2.1. Default settings were used to perform the hybrid scaffolding. Dual enzyme hybrid scaffolding (incorporating both DLS and NLRS maps) resulted in a hybrid assembly with N50 of 13.7 Mb (scaffold only N50 = 34.28 Mb) for a total length of 1,414.64 Mb and consisted of 154 scaffolds (999.96 Mb) and 6,336 unscaffolded contigs (414.68 Mb, Table 1).

### Anchoring

The first round of analyses on the whole hybrid assembly with ALLMAPS identified 31 scaffolds that were sufficient to represent one haploid genome. Seven pseudomolecules (chr8, 9, 12, 13, 14, 16, and 17) were covered by one scaffold each. For each one of the remaining pseudomolecules (covered by two to three scaffolds), overlapping regions between scaffolds were searched and eight overlapping regions were removed manually. When re-anchored using ALLMAPS, a primary haploid genome (MDGGph_v1.0) assembly spanning 719 Mb in 17 chromosome-scale pseudomolecules (chr1-17) was generated. Its size is in agreement with the estimated genome size of 710 Mb. MDGGph_v1.0 served as haploid reference assembly in order to map long and short reads generating the graph-based genome for subsequent k-mer analysis. The remaining sequences were assembled again using ALLMAPS into a second set of 17 pseudomolecules (secondary haploid genome assembly MDGGsh_v1.0, chr_s1-17) spanning 601 Mb, with 80 Mb unanchored sequences. In four cases the secondary haploid assembly showed longer pseudomolecules when compared to the primary assembly (chromosomes chr_s5, chr_s 6, chr_s11 and chr_s 14, supplementary Table 1). This is possibly due to differences in chromosome length or misassembles. Unplaced sequences (n = 2,853) spanned only 5.7% of the total assembly (Table 1).

Genome visualization displayed Illumina read coverage, SNP phase, and correlation of genetic and physical order for each chromosome pair on the assembly, as well as links between both alleles of each SNP marker (Genome visualization, supplementary data). With the exception of a region on chr_s10, all collapsed regions (with an Illumina read coverage of about 130×) were present in MDGGph_v1.0. Our assembly show collinearity with the genetic map of ‘Fuji’ × ‘Gala’. Phase switches were observed along the pseudomolecules and, as expected, much lower SNP phase information was found in these collapsed regions (see Genome visualization, supplementary data).

The complete diploid assembly of ‘Gala Galaxy’ (MDGGdi_v1.0) obtained by combining primary and secondary haploid assemblies is available in Genbank as Bioproject XXXXXX. Genome assembly statistics are shown in Table 1, chromosome sizes are shown in supplementary Table 1, while analysis results of BUSCOs identified in MDGGdiv1.0, MDGGph_V1.0 and GDDH13 are shown in supplementary Table 2. The number of complete BUSCOs identified in MDGGdi_v1.0 was 1,387 (96.3%), of which 467 (32.4%) were present as single-copy and 920 (63.9%) as duplicated (Table 3). The same analysis on the primary haploid assembly MDGGph_v1.0 identified 1,347 (93.6%) complete BUSCOs, with 943 (65.5%) being single-copy and 404 (28.1%) were duplicated (supplementary Table 2).

### Gene annotation

Using RNAseq data, a total of 74,900 leaf transcripts encoded by 41,264 genes were identified and functionally annotated as described in Knorst et al. (2019). Of the 74,900 proteins searched, 64,765 had an annotation assigned, of which 59,039 were assigned at least one GO term. The *de novo* annotated primary assembly MDGGph_v1.0 and the GDDH13 assembly were used to generate a synthenic dotplot, confirming a high collinearity between the two assemblies (Supplementary Figure 1).

### K-mer analysis

Kat spectra plots were used to investigate k-mer multiplicity distribution and counting the occurrence of each k-mer in the investigated assemblies. The distribution of the k-mer counts in the KAT spectra plot revealed two peaks: the first peak at ×=27 represents the heterozygous content (and hence unique sequences) and the second peak at ×=58 the homozygous (Figure 1). The color of the area under the curve indicated how often k-mers were found in the investigated reference assemblies: black corresponds to k-mers missing in the assembly, while red and violet correspond to k-mers counted one and twice, respectively. The black area in the spectrum generated using GDDH13 is clearly larger than the one generated using MDGGdi_v1.0 (Figure 1A and 1B). The spectrum generated using the phased graph-based assembly (based on MDGGph_v1.0) shows a large black area, but also a violet area under the peak for heterozygous content (Figure 1C).

**Figure 1:**
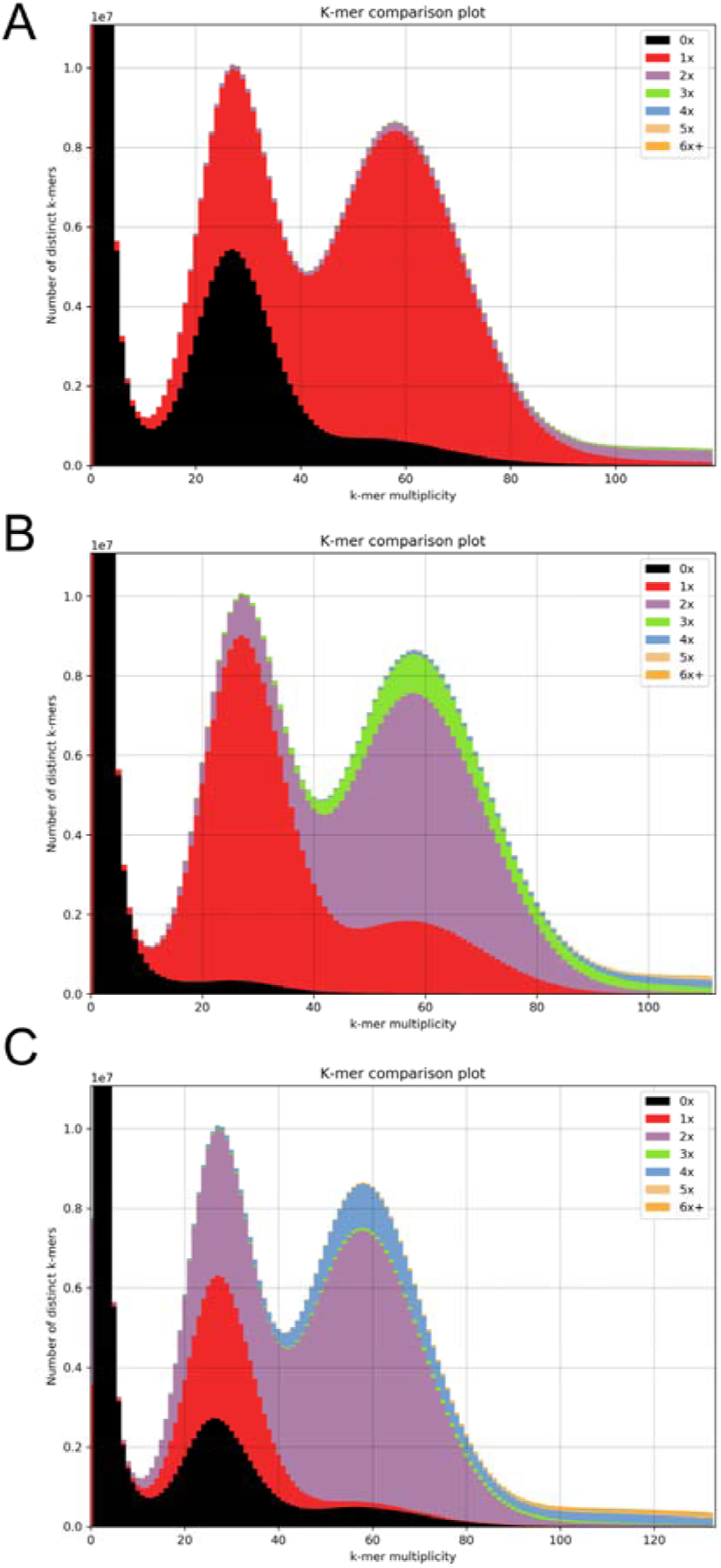
K-mers spectra generated using k-mer analysis toolkit (23-mers) with the Illumina reads of ‘Gala Galaxy’ against the double-haploid GDDH13 reference genome (A), the diploid genome reference MDGGdi_v1.0 (B) and the graph-based diploid phased assembly generated with WhatsHap (C). The overall curve shape indicates the frequency of the 23-mers in the original Illumina data (apple cv. ‘Gala Galaxy’), while the colored area indicate if the 23-mers are found once (red), twice (violet) or more often (other colors) in the reference genome. The black area indicate reads that are present in the Illumina sequencing data but absent in reference assemblies.

## Discussion

Here, we present a diploid assembly of the heterozygous apple cultivar ‘Gala Galaxy’ (MDGGdi_v1.0). Optical mapping was essential in order to increase the contiguity of the assembly, as shown by the 60-fold increase of N50 values, reaching 34.28 Mb for the hybrid scaffolds only (Table 1) and corresponding to a two-fold increase compared to the GDDH13 (Daccord, et al. 2017) and HFTH1 (Zhang, et al. 2019) apple genome assemblies. Despite being diploid, this assembly is unphased, as phase switches assessed by SNP markers were observed (Supplementary data). Phase switches are probably due to the unzip approach in FALCON, where the choice of which variant to include in the primary contigs is based on length – resulting in chimeric contigs. Due to the Mb resolution of the optical map, this data was not helpful in rearranging chimeric scaffolds into correctly phased scaffolds. BUSCO analyses confirmed that the assembly MDGGdi_v1.0 is complete, with only 3.7% missing BUSCOs. The diploid assembly presented in this study represents an advancement over existing resources. Further improvements might be achieved by separating the collapsed haplotypes and resolving the phase switches in to a correctly phased genome. Moreover, gene annotation could further be improved by integrating transcriptome data from different tissue types (e.g. fruits or flowers).

Our results clearly show that using a haploid reference is suboptimal to analyse genomic data derived from the heterozygous apple and might lead to sequence information loss, as indicated by the black area in Figure 1A. This occurred despite the closeness of these two genotypes (as ‘Gala’ is an offspring of ‘Golden Delicious’, from which the doubled-haploid GDDH13 was derived). Furthermore, the k-mer analysis indicated that a graph-based approach generated an assembly in which some allelic regions were absent while other were artificially duplicated (black and violet area in Figure 1C, respectively. Thus, by running WhatsHap on a haploid reference, the second allele could not be recovered as well as the *de novo* assembly by FALCON Unzip did. This may be a consequence of a high divergence between allelic regions and therefore, haplotype homology was not resolved correctly by this approach.

This diploid genome will thus be the appropriate reference for achieving the most information from resequencing projects of sport mutants of this important apple cultivar. Understanding the genome organization and including genomic information in the breeding process will enable the rapid development of resilient cultivars allowing a more sustainable production of this important fruit.

## Supporting information

Genome Visualization

## Availability of supporting data

The Illumina sequencing reads of each sequencing library and the RNA-seq data have been deposited at SRA with the accessions numbers XXXXXXX. The genome assembly MDGGdi_v1.0 was deposited at NCBI as Bioproject XXXXXXXX. The genome visualization is available as supplementary data.

## Authors’ contributions

GALB, BS, and AP designed the study. GALB and GR assembled the genome. IS and GALB collected leaf samples and extracted DNA. GALB, GR, SAY, and DC analyzed the data. All authors participated to the writing of the manuscript and approved the final version.

## Acknowledgments

We acknowledge the Functional Genomic Centre Zurich, Switzerland, especially Dr. Lucy Poveda, Andrea Patrignani and Dr. Catharine Aquino-Fournier for the support in generating the genome sequencing data, the Fruit-Growing Extension Group of Agroscope, Switzerland, for taking care of the plant material and the Method Development and Analytics Group of Agroscope, Switzerland, for sharing laboratory infrastructures. Part of the analyses were performed on the CyVerse cyberinfrastructure, which was supported by the National Science Foundation under Award Numbers DBI-0735191, DBI-1265383, and DBI-1743442. URL: www.cyverse.org

