## Supplementary material for "Chromosome-scale *de novo* diploid assembly of the apple cultivar ‘Gala Galaxy’": Genome Visualization

Chr1.pdf

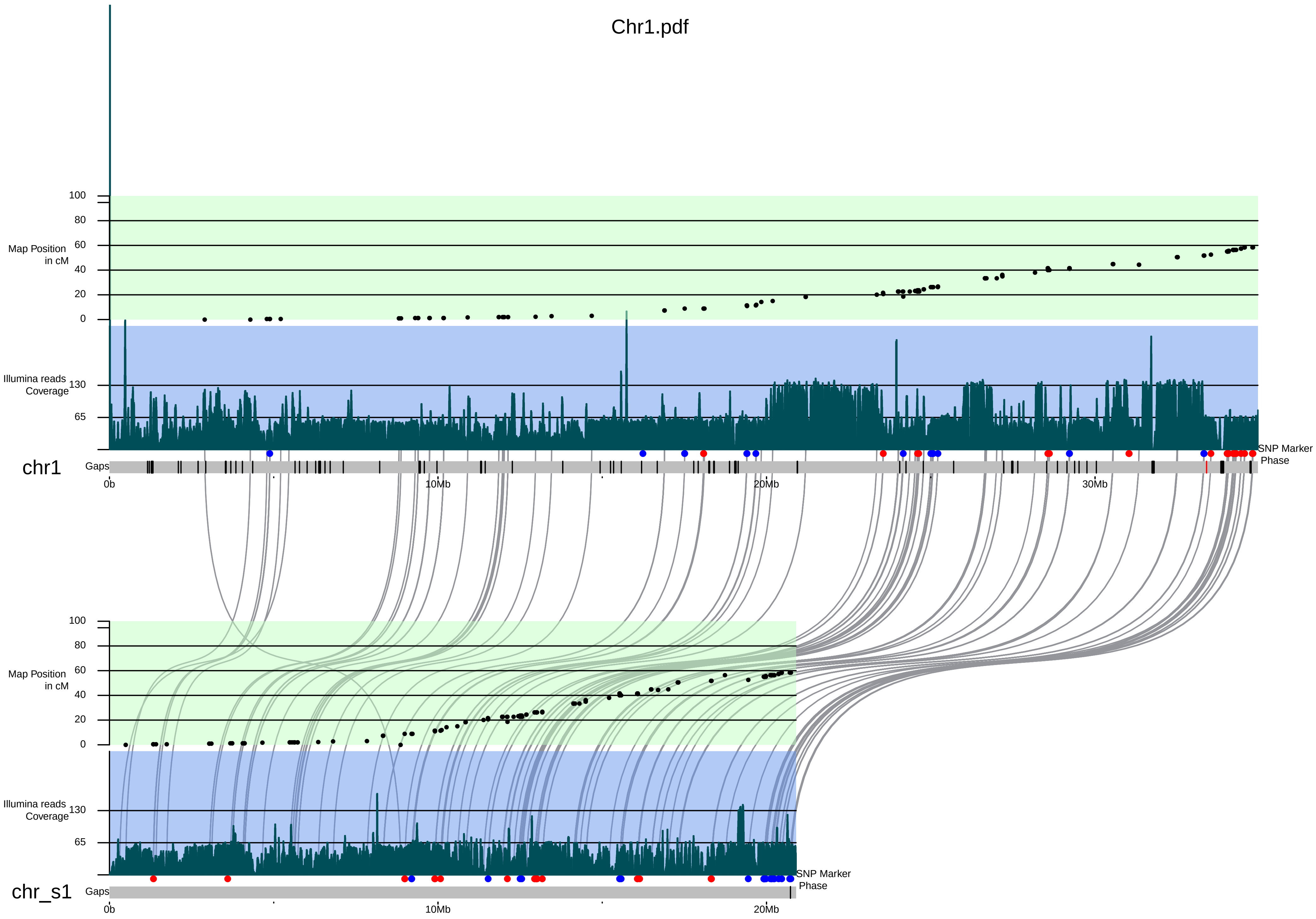

100
80
60
40
20
0
Map Position
in cM
Illumina reads
Coverage
130
65
chr1
Gaps
SNP Marker
Phase
0b
10Mb
20Mb
30Mb
100
80
60
40
20
0
Map Position
in cM
Illumina reads
Coverage
130
65
chr_s1
Gaps
SNP Marker
Phase
0b
10Mb
20Mb

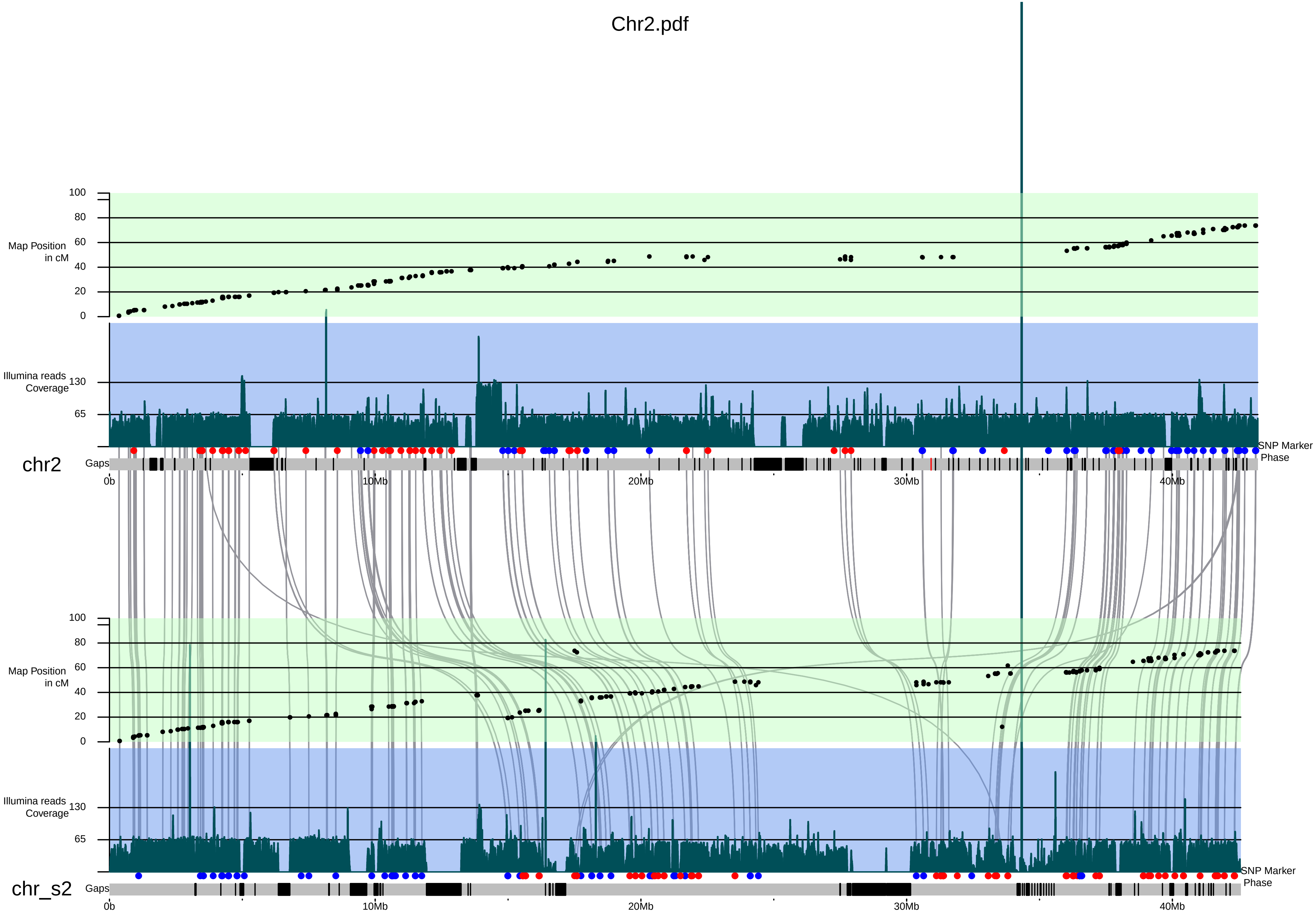
Chr2.pdf
100
80
60
40
20
0
Map Position
in cM
Illumina reads
Coverage
130
65
chr2
Gaps
SNP Marker
Phase
0b
10Mb
20Mb
30Mb
40Mb
100
80
60
40
20
0
Map Position
in cM
Illumina reads
Coverage
130
65
chr_s2
Gaps
SNP Marker
Phase
0b
10Mb
20Mb
30Mb
40Mb

### Chr3.pdf

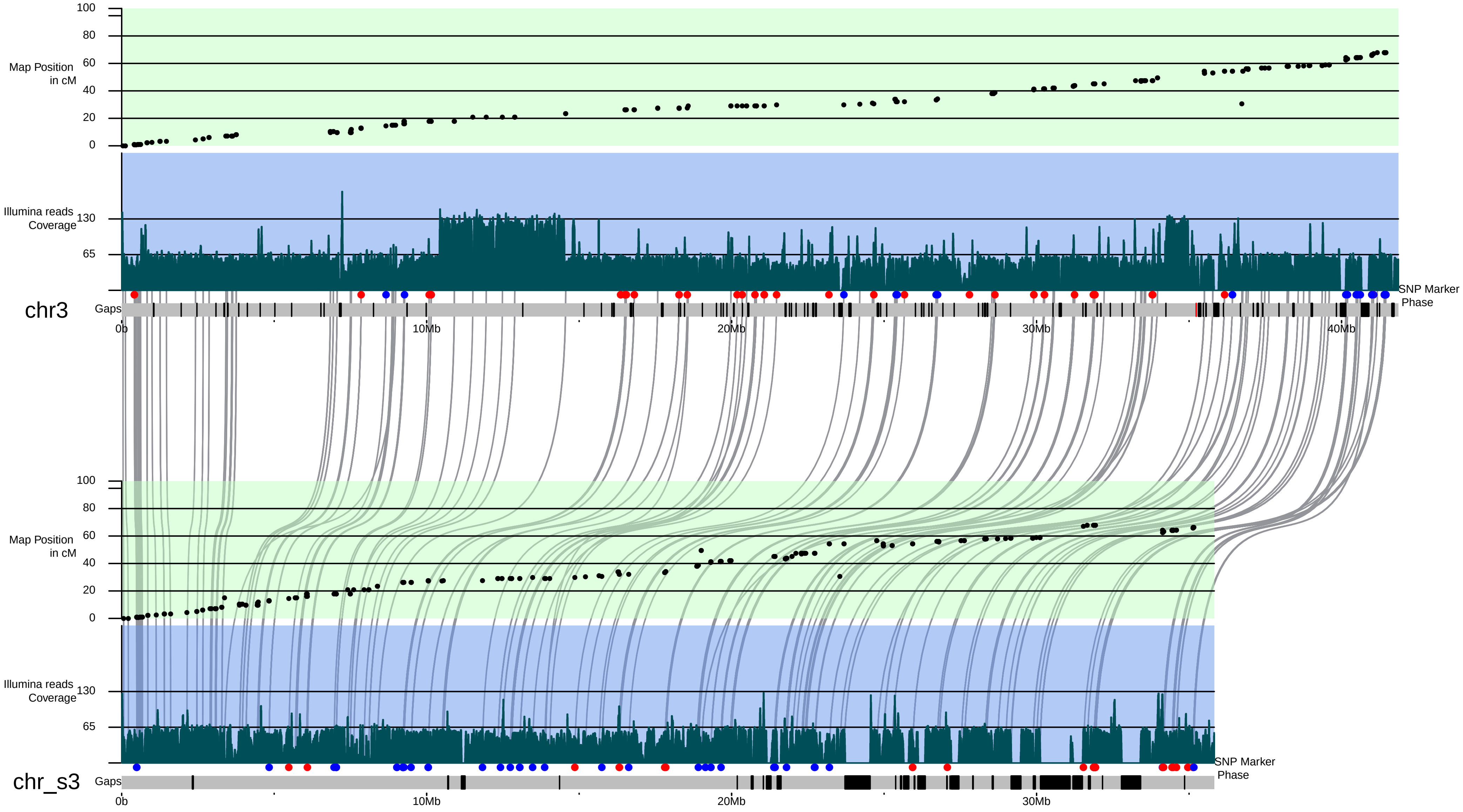

Map Position in cM
100
80
60
40
20
0
Illumina reads Coverage
130
65
chr3
Gaps
SNP Marker Phase
0b
10Mb
20Mb
30Mb
40Mb
Map Position in cM
Illumina reads Coverage
chr_s3
Gaps
SNP Marker Phase

Chr4.pdf

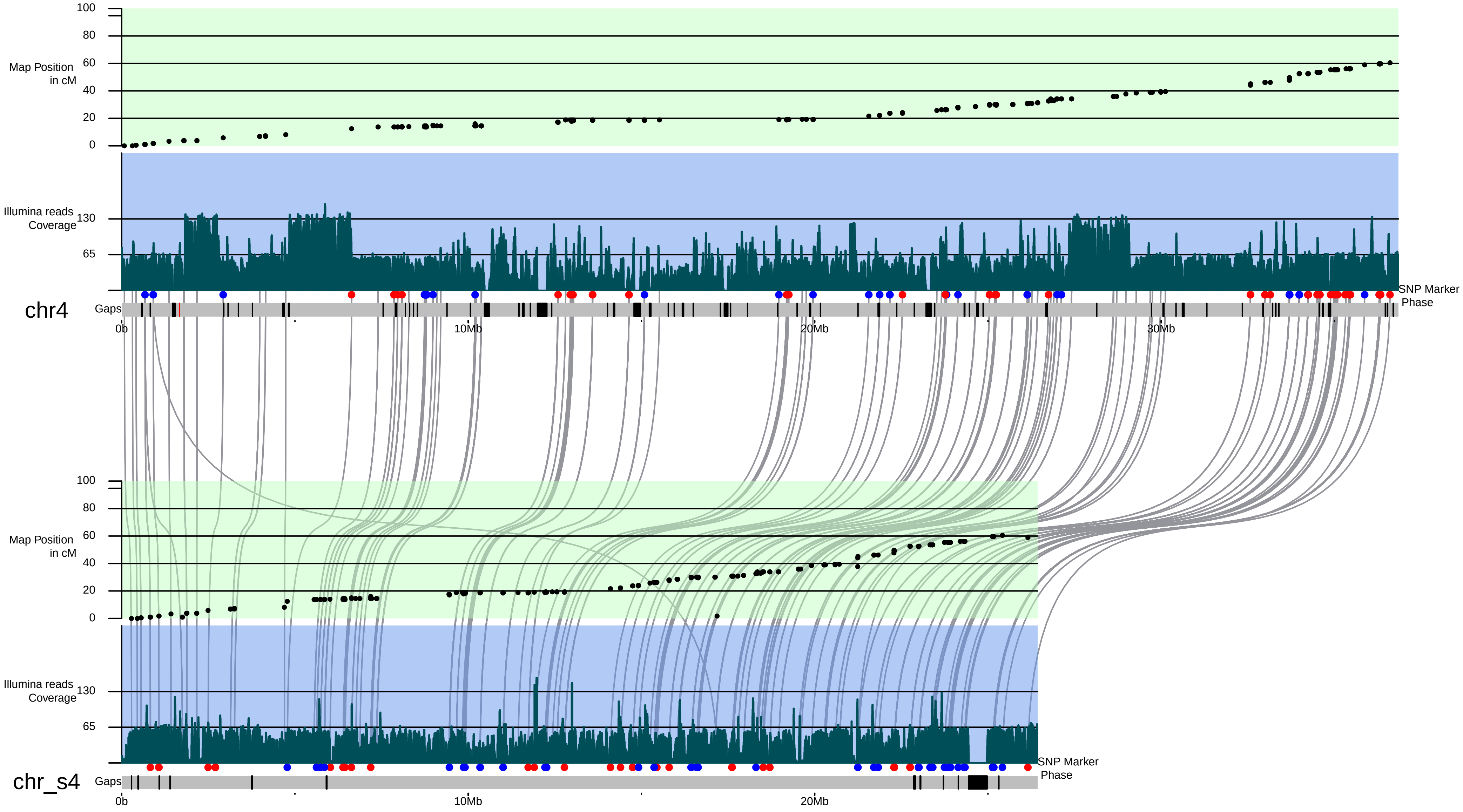

100
80
60
40
20
0
Map Position in cM
Illumina reads Coverage
130
65
SNP Marker Phase
chr4
Gaps
0b
10Mb
20Mb
30Mb
chr_s4

### Chr5.pdf

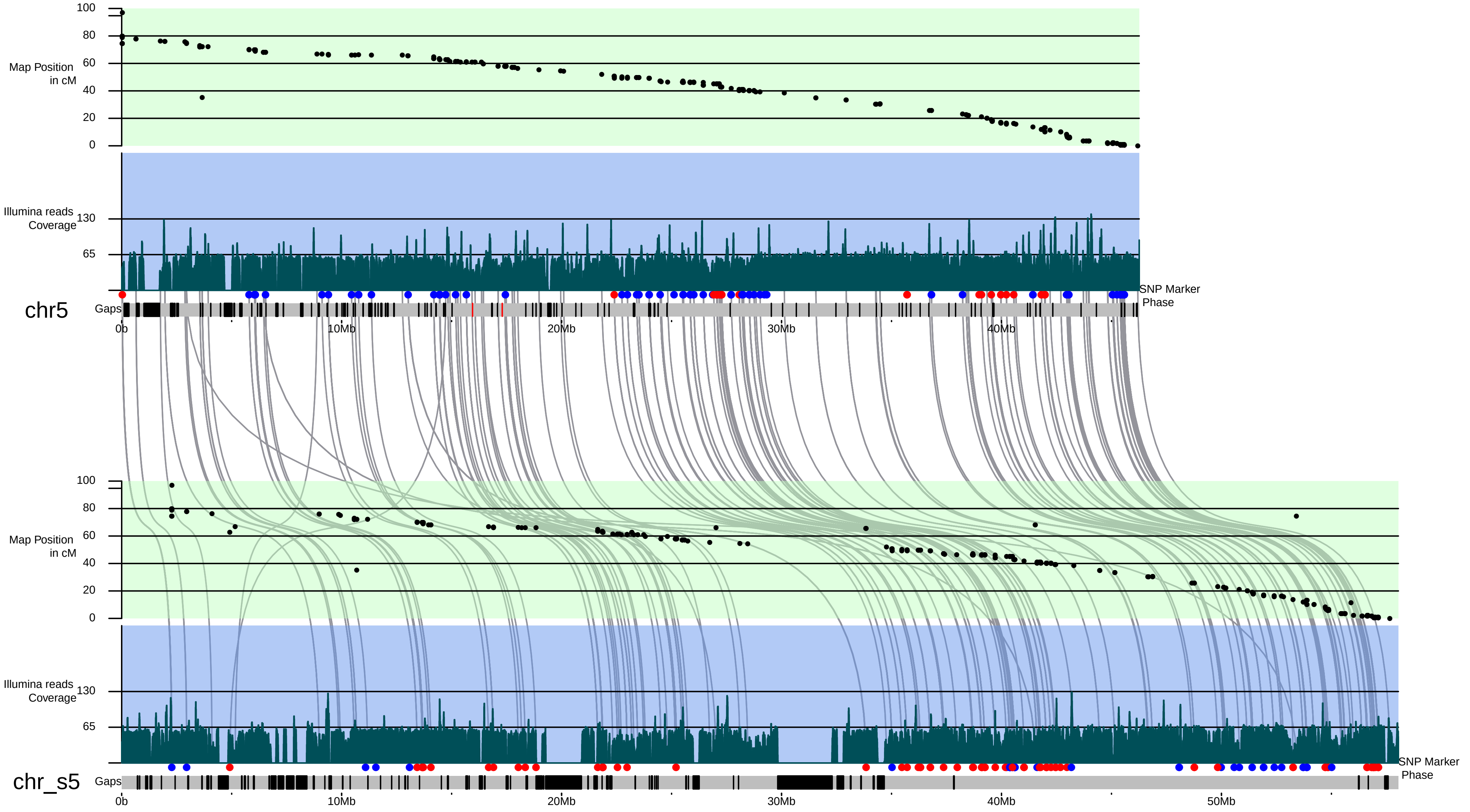

Map Position in cM
100
80
60
40
20
0
Illumina reads Coverage
130
65
SNP Marker Phase
chr5
Gaps
0b
10Mb
20Mb
30Mb
40Mb
Map Position in cM
100
80
60
40
20
0
Illumina reads Coverage
130
65
SNP Marker Phase
chr_s5
Gaps
0b
10Mb
20Mb
30Mb
40Mb
50Mb

### Chr6.pdf

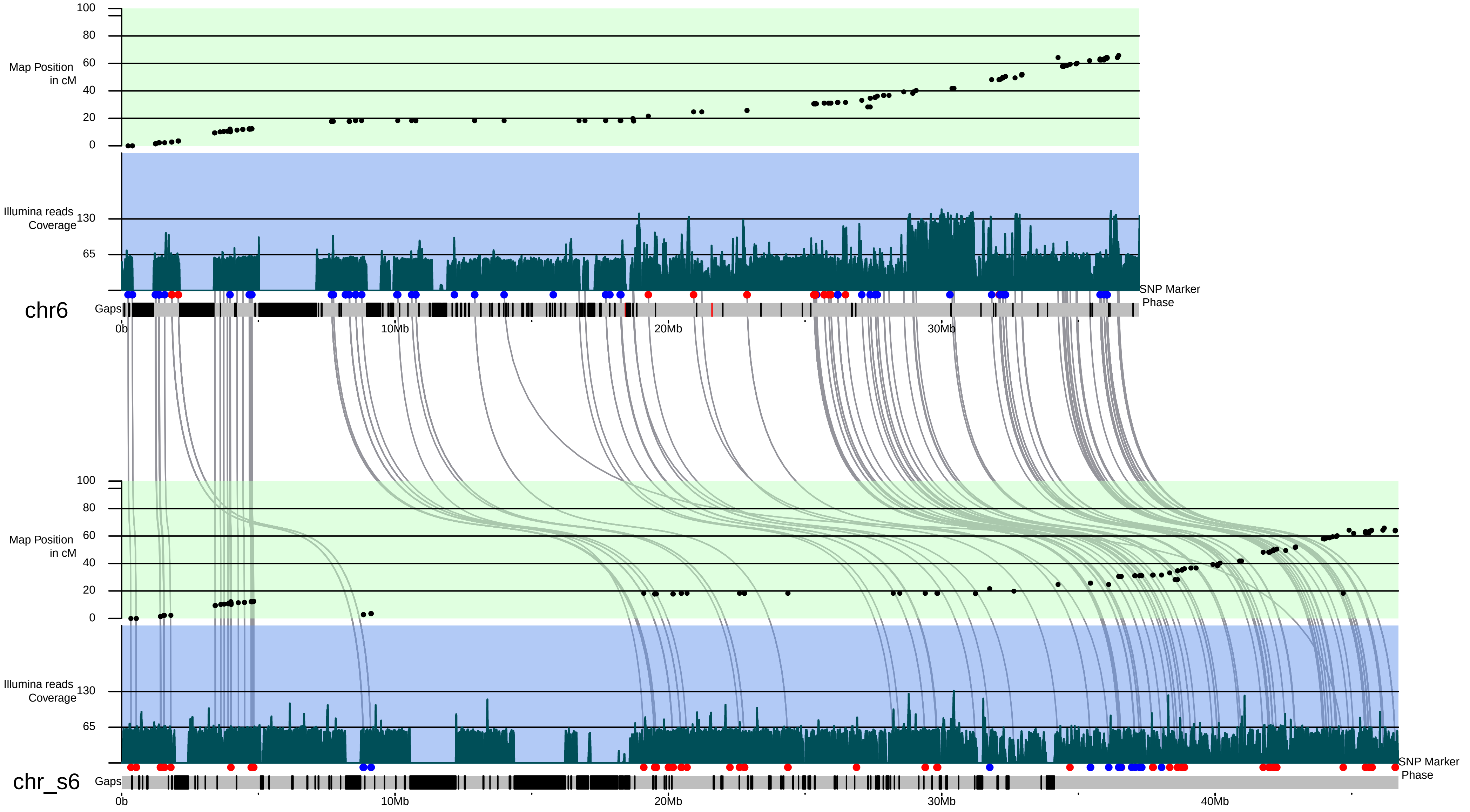
Map Position in cM
100
80
60
40
20
0
Illumina reads Coverage
130
65
SNP Marker Phase
chr6
Gaps
0b
10Mb
20Mb
30Mb
chr_s6
40Mb

### Chr7.pdf

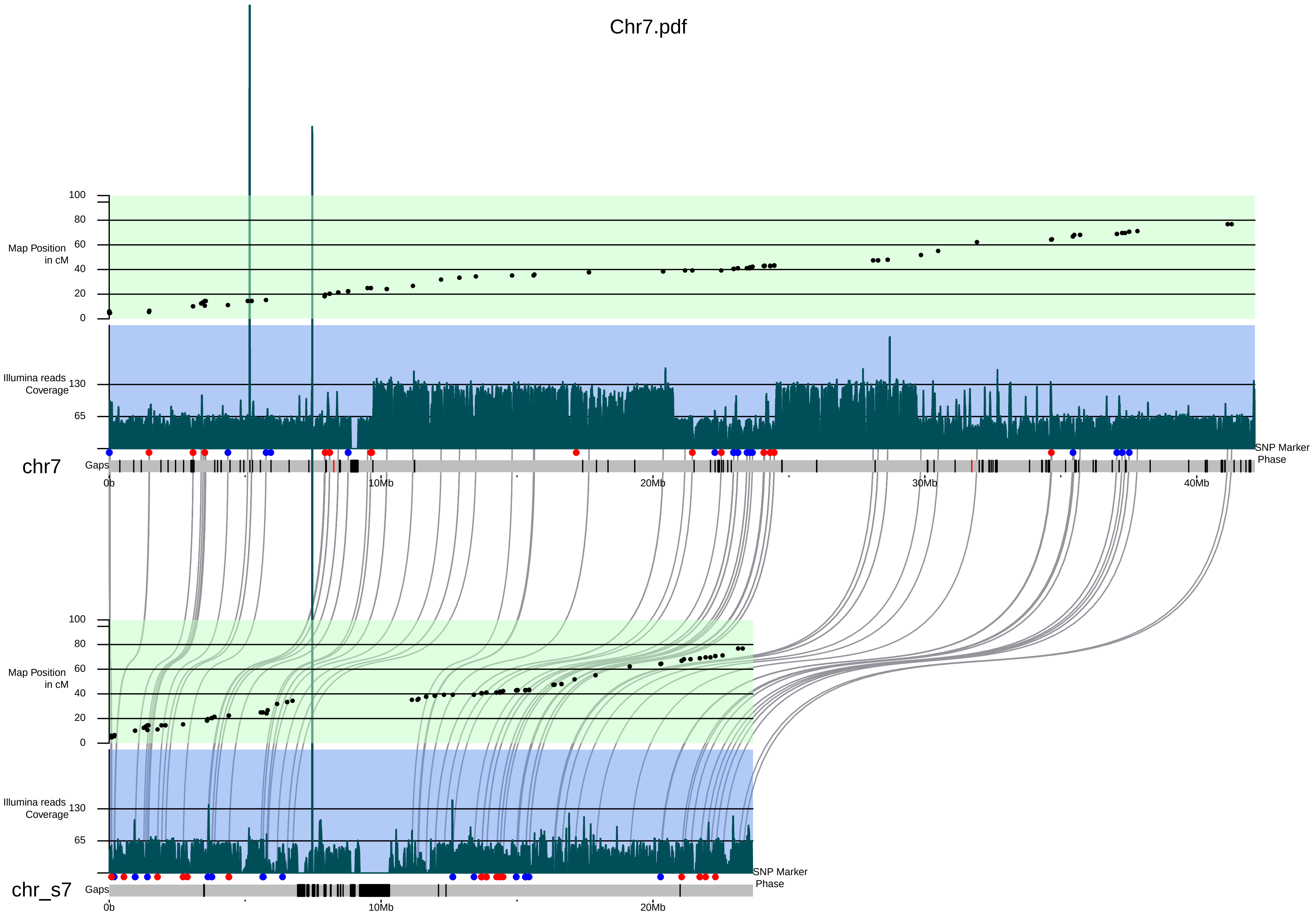

100
80
60
40
20
0
Map Position
in cM
Illumina reads
Coverage
130
65
chr7
Gaps
SNP Marker
Phase
0b
10Mb
20Mb
30Mb
40Mb
100
80
60
40
20
0
Map Position
in cM
Illumina reads
Coverage
130
65
chr_s7
Gaps
SNP Marker
Phase
0b
10Mb
20Mb

Chr8.pdf

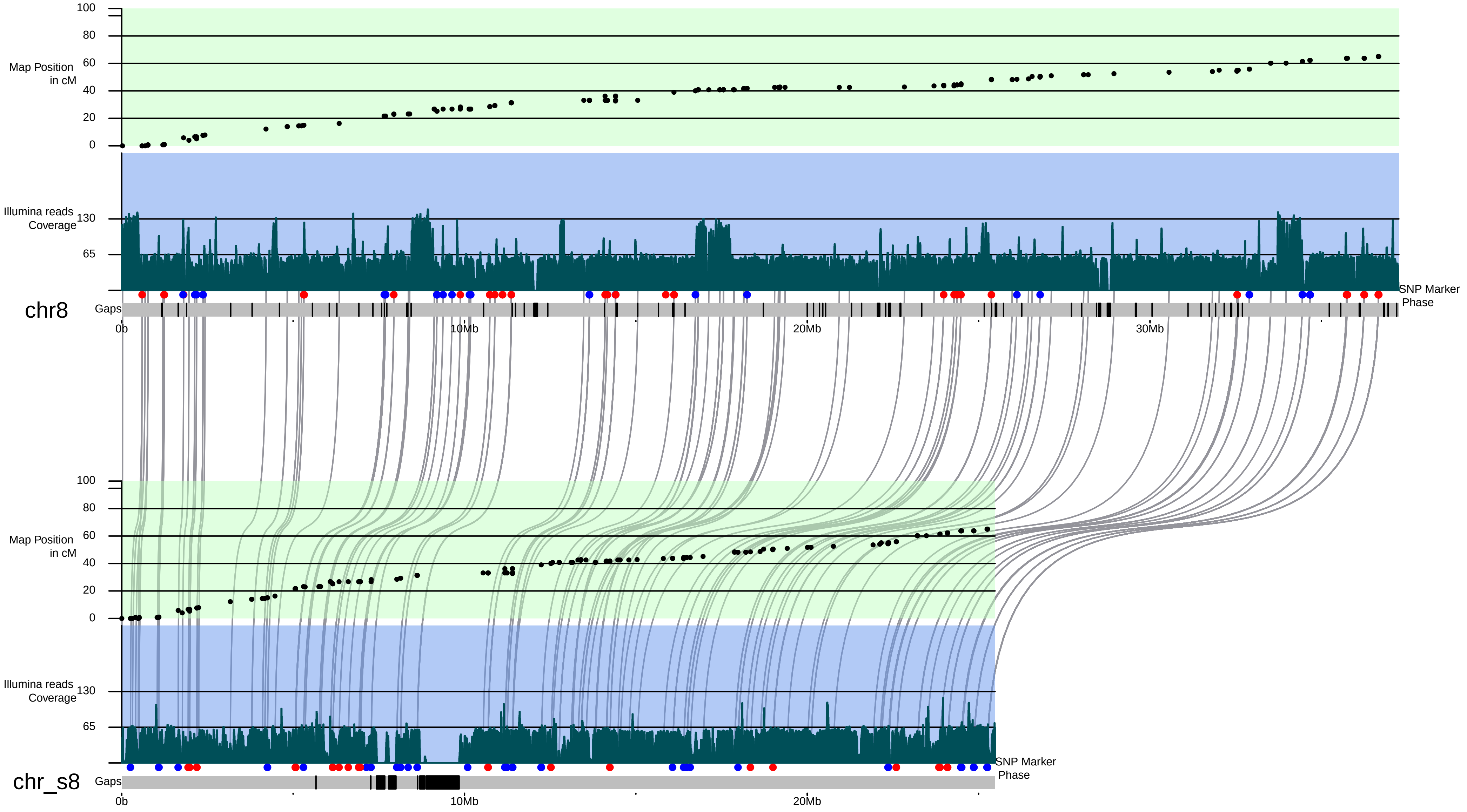

Map Position in cM
100
80
60
40
20
0
Illumina reads Coverage
130
65
SNP Marker Phase
chr8
Gaps
0b
10Mb
20Mb
30Mb
Map Position in cM
100
80
60
40
20
0
Illumina reads Coverage
130
65
SNP Marker Phase
chr_s8
Gaps
0b
10Mb
20Mb

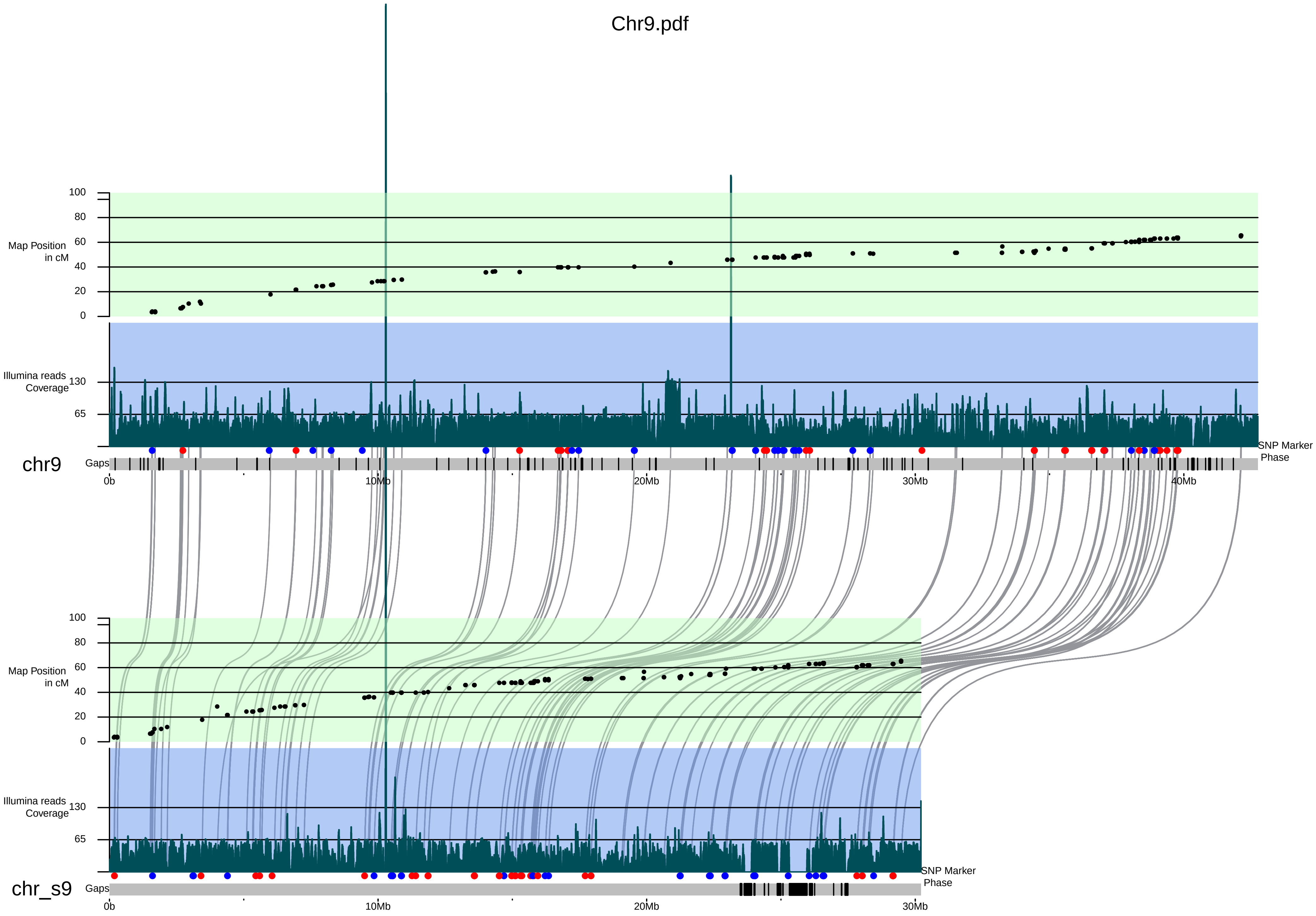

Chr9.pdf
Map Position in cM
100
80
60
40
20
0
Illumina reads Coverage
130
65
chr9
Gaps
SNP Marker Phase
0b
10Mb
20Mb
30Mb
40Mb
Map Position in cM
100
80
60
40
20
0
Illumina reads Coverage
130
65
chr_s9
Gaps
SNP Marker Phase
0b
10Mb
20Mb
30Mb

Chr10.pdf

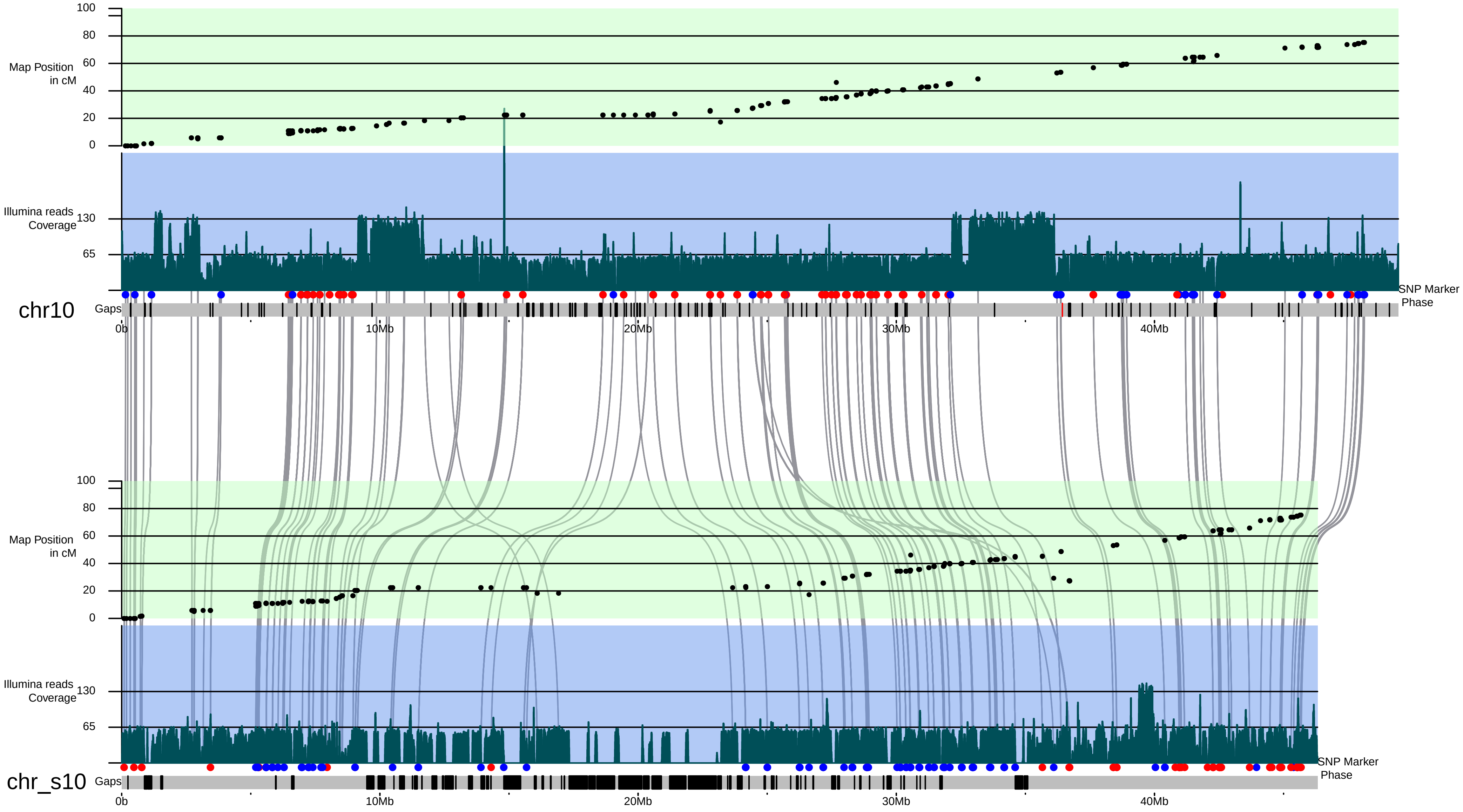

Map Position in cM
100
80
60
40
20
0
Illumina reads Coverage
130
65
SNP Marker Phase
chr10
Gaps
0b
10Mb
20Mb
30Mb
40Mb
Map Position in cM
100
80
60
40
20
0
Illumina reads Coverage
130
65
SNP Marker Phase
chr_s10
Gaps
0b
10Mb
20Mb
30Mb
40Mb

### Chr11.pdf

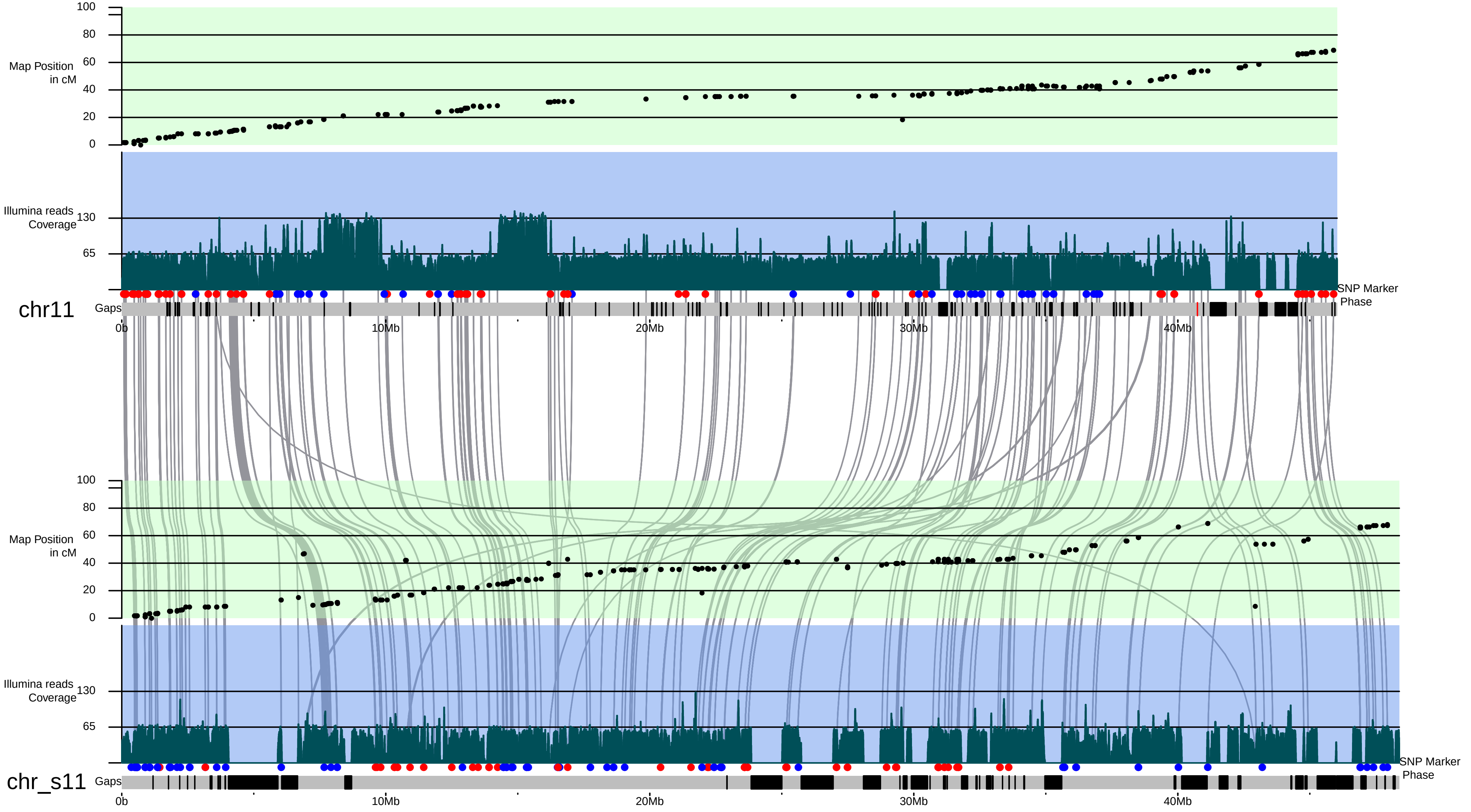

Map Position in cM
100
80
60
40
20
0
Illumina reads Coverage
130
65
chr11
Gaps
SNP Marker Phase
0b
10Mb
20Mb
30Mb
40Mb
Map Position in cM
100
80
60
40
20
0
Illumina reads Coverage
130
65
chr_s11
Gaps
SNP Marker Phase
0b
10Mb
20Mb
30Mb
40Mb

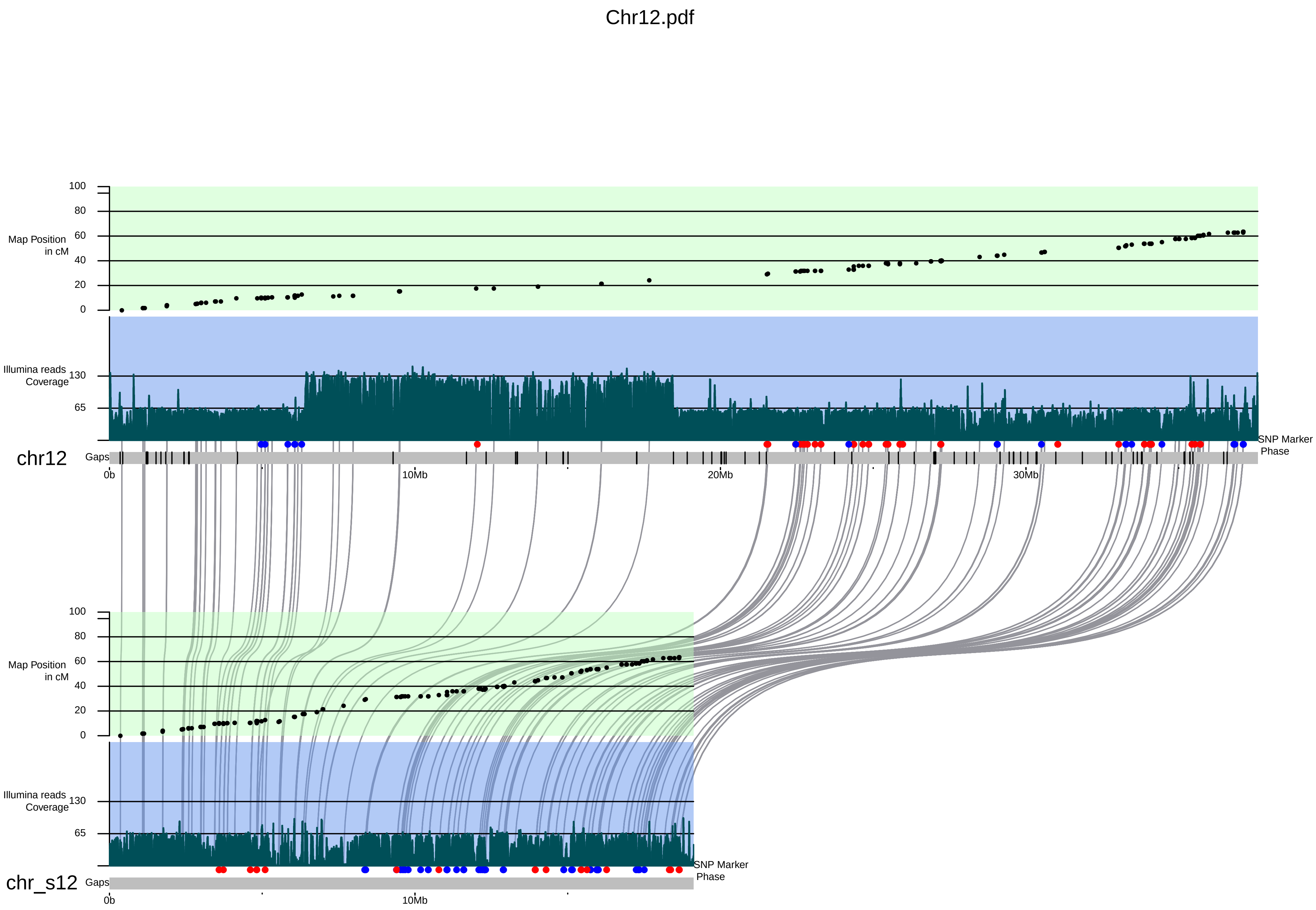
Chr12.pdf
Map Position in cM
100
80
60
40
20
0
Illumina reads Coverage
130
65
chr12
Gaps
0b
10Mb
20Mb
30Mb
SNP Marker Phase
chr_s12

Chr13.pdf

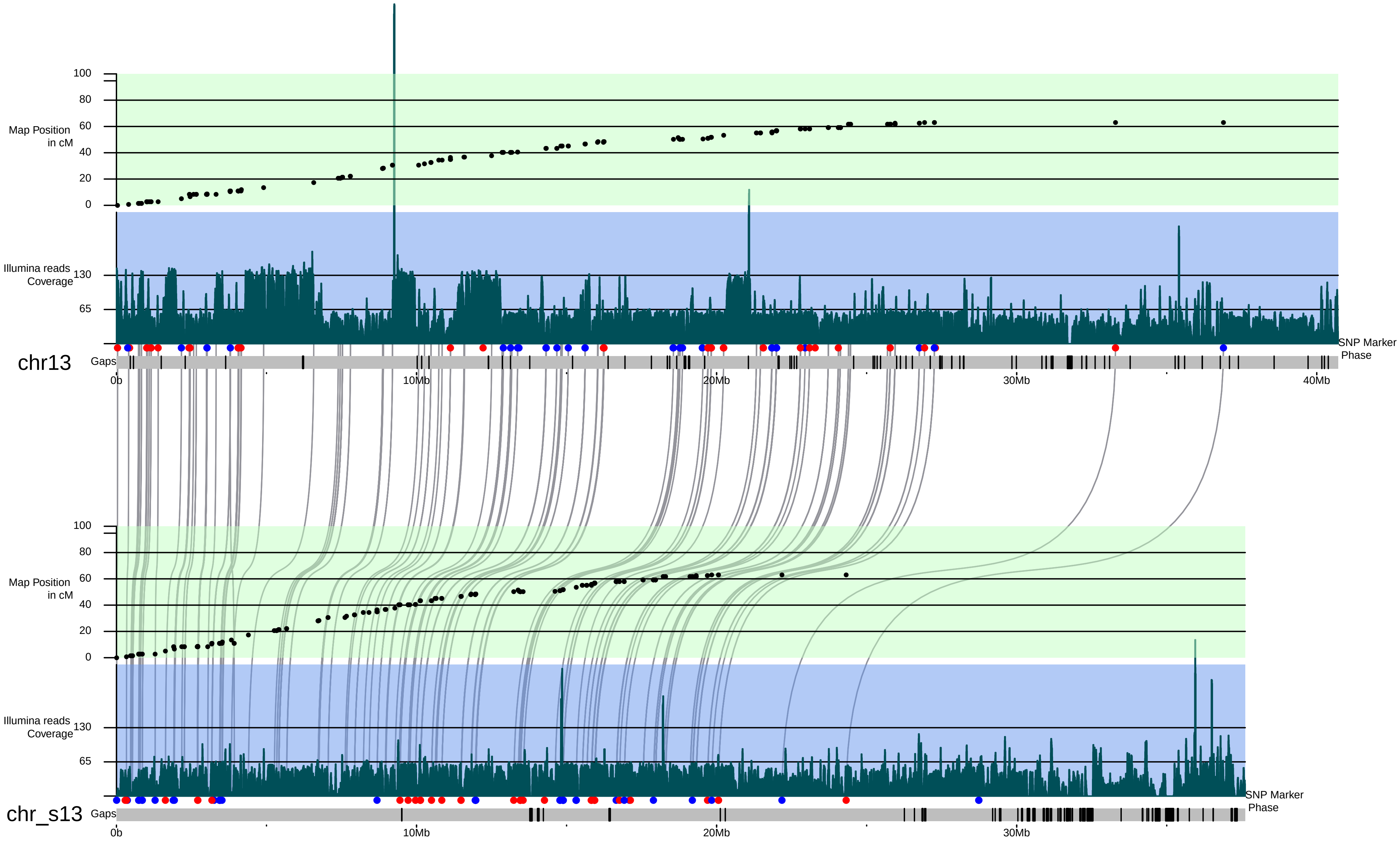

100
80
60
40
20
0
Map Position in cM
Illumina reads Coverage
130
65
chr13
Gaps
SNP Marker Phase
0b
10Mb
20Mb
30Mb
40Mb
100
80
60
40
20
0
Map Position in cM
Illumina reads Coverage
130
65
chr_s13
Gaps
SNP Marker Phase
0b
10Mb
20Mb
30Mb

### Chr14.pdf

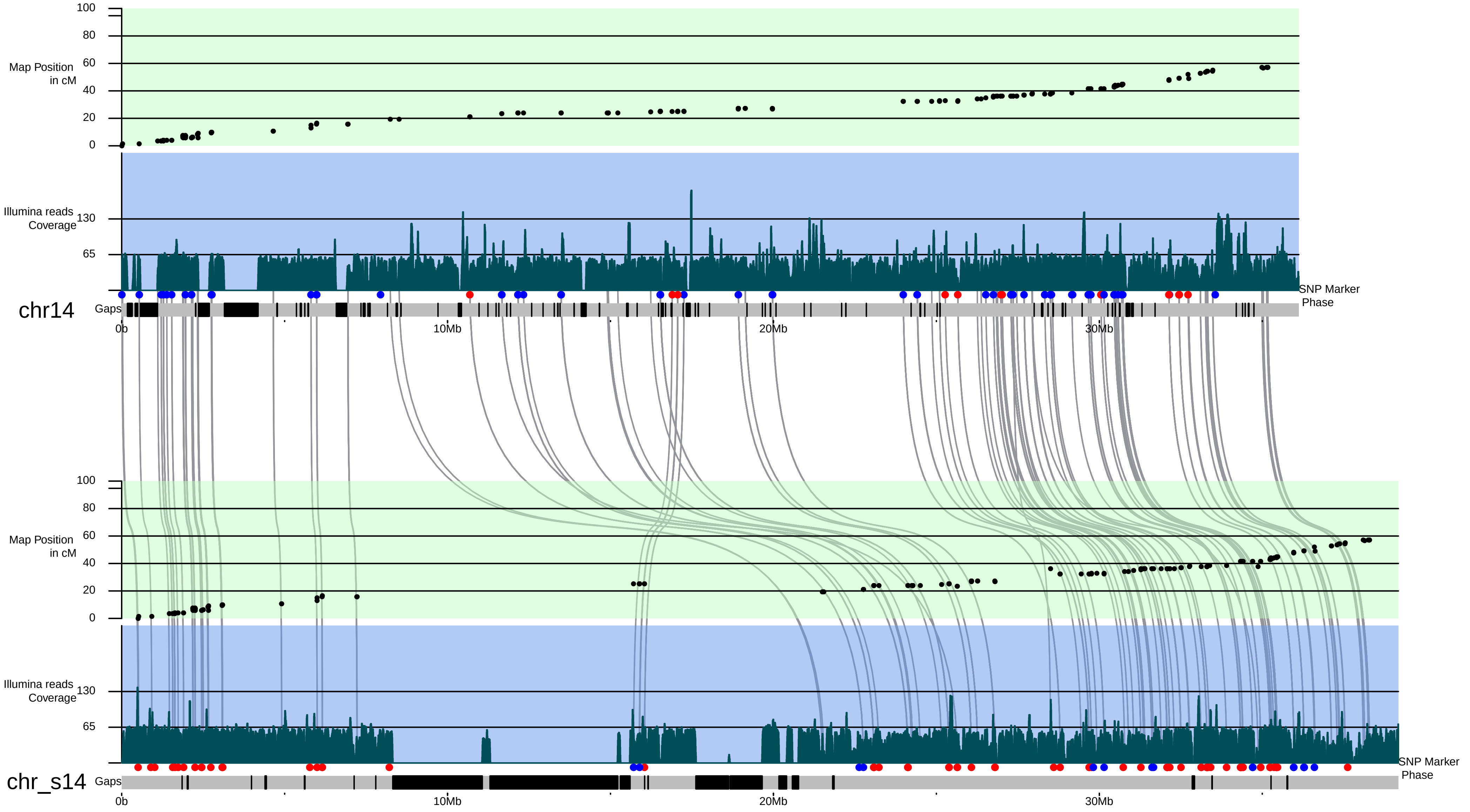

Map Position in cM
100
80
60
40
20
0
Illumina reads Coverage
130
65
SNP Marker Phase
chr14
Gaps
0b
10Mb
20Mb
30Mb
Map Position in cM
100
80
60
40
20
0
Illumina reads Coverage
130
65
SNP Marker Phase
chr_s14
Gaps
0b
10Mb
20Mb
30Mb

Chr15.pdf

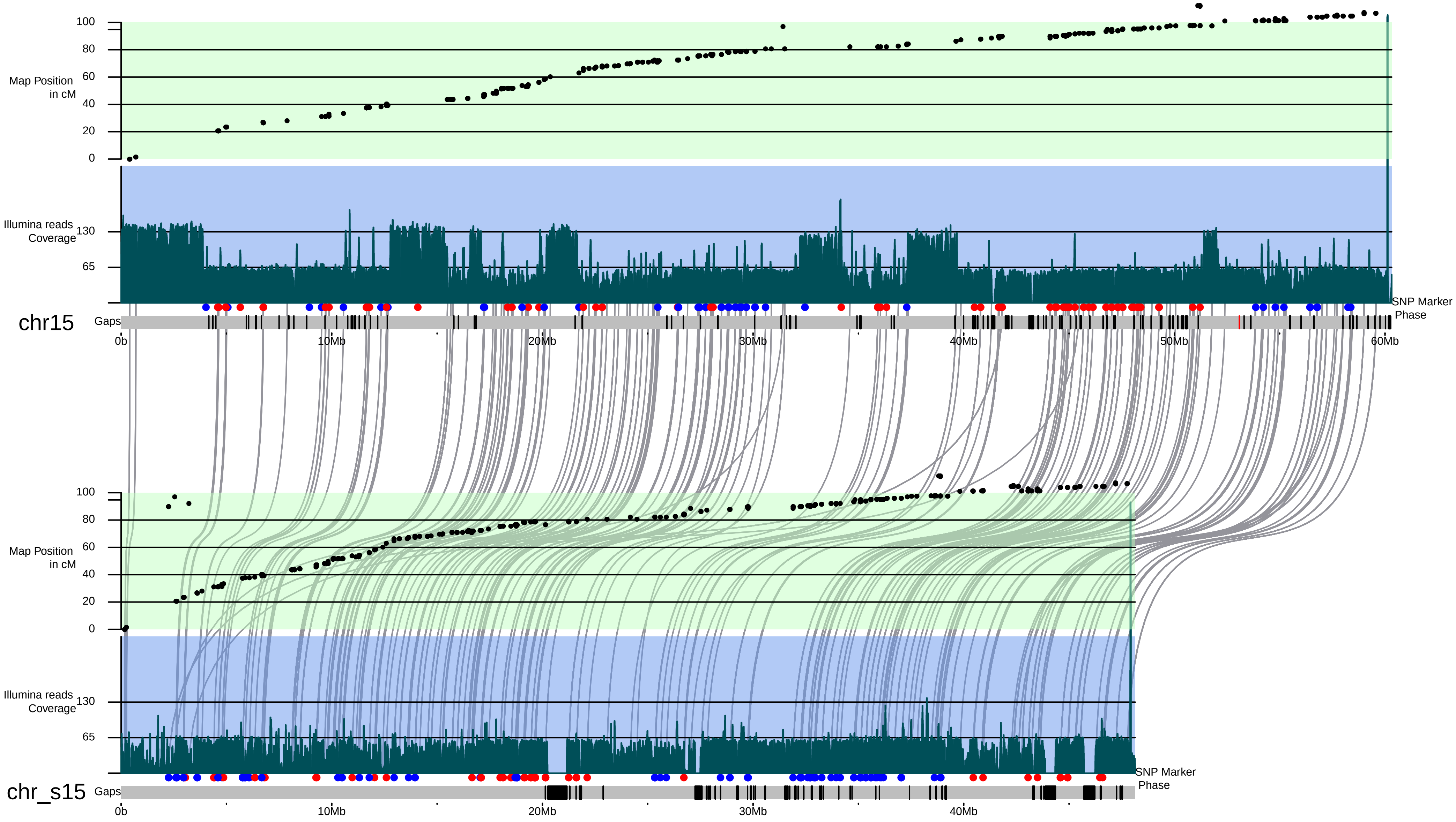

Map Position in cM
Illumina reads Coverage
chr15
Gaps
SNP Marker Phase
0b
10Mb
20Mb
30Mb
40Mb
50Mb
60Mb
chr_s15

### Chr16.pdf

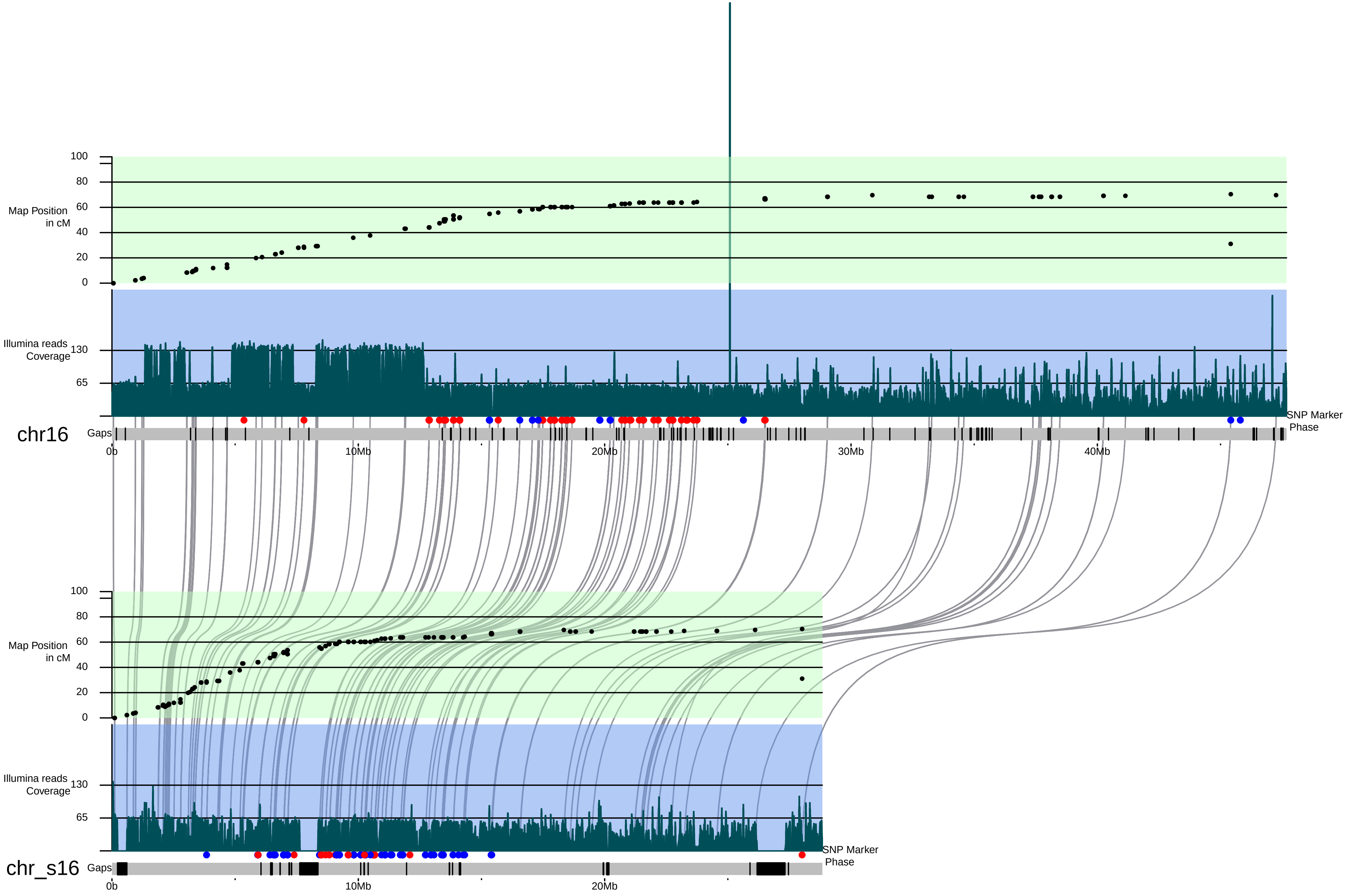

Map Position in cM
100
80
60
40
20
0
Illumina reads Coverage
130
65
chr16
Gaps
SNP Marker Phase
0b
10Mb
20Mb
30Mb
40Mb
Map Position in cM
100
80
60
40
20
0
Illumina reads Coverage
130
65
chr_s16
Gaps
SNP Marker Phase
0b
10Mb
20Mb

### Chr17.pdf

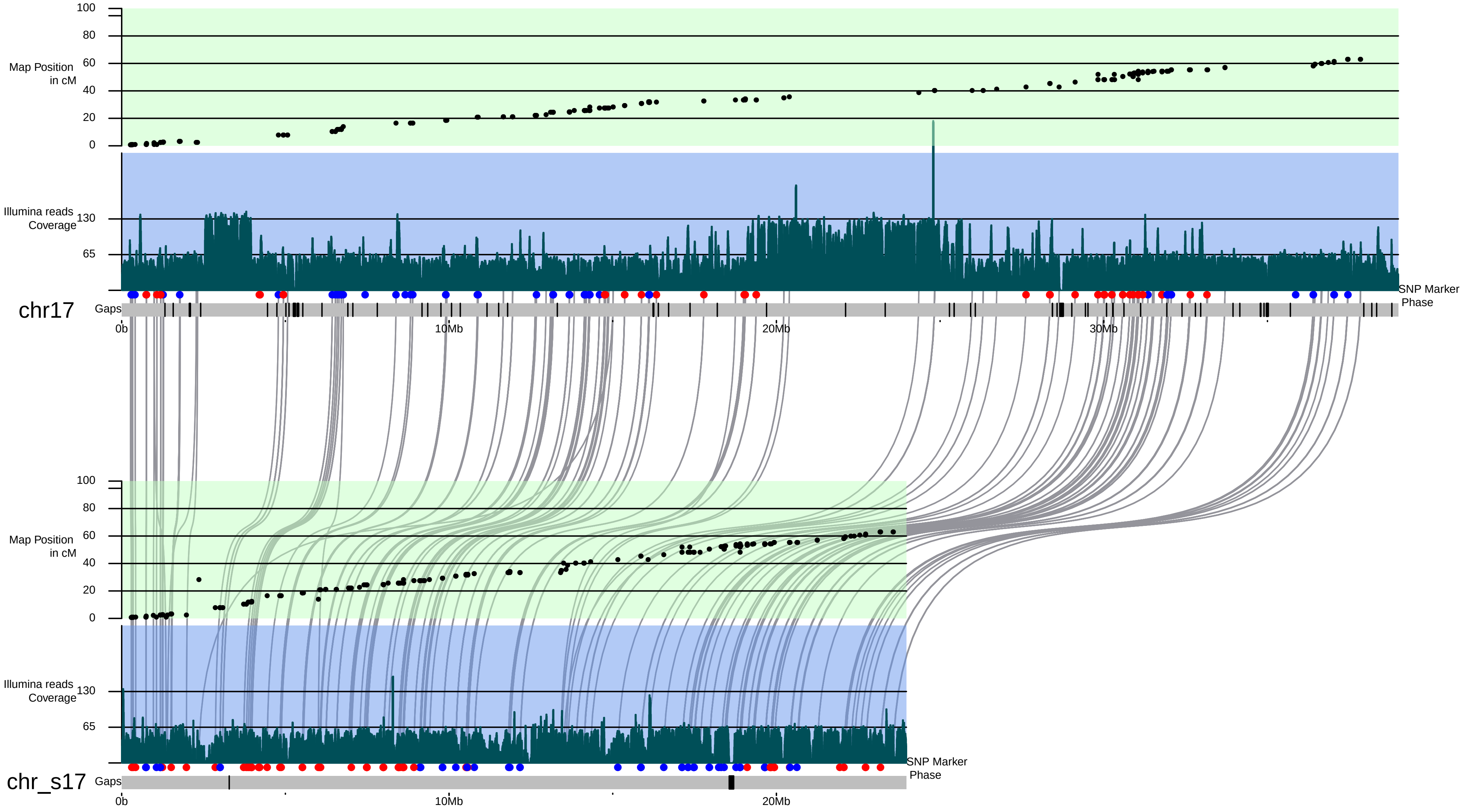

100
80
60
40
20
0
Map Position
in cM
Illumina reads
Coverage
130
65
SNP Marker
Phase
chr17
Gaps
0b
10Mb
20Mb
30Mb
100
80
60
40
20
0
Map Position
in cM
Illumina reads
Coverage
130
65
SNP Marker
Phase
chr_s17
Gaps
0b
10Mb
20Mb
