## Supplementary Methods for "Chromosome-scale *de novo* diploid assembly of the apple cultivar ‘Gala Galaxy’"

**DNA extraction and library preparation.**

Genomic DNA was isolated from two grams of field-grown apple leaves according to the PacBio “Preparing Arabidopsis Genomic DNA for Size-Selected ~20 kb SMRTbell Libraries” protocol. Quantification and quality assessment was made using the Qubit Fluorometer dsDNA Broad Range assay (Thermo Fisher Scientific, Waltham, MA, USA) and the Bioanalyzer 2100 12K DNA Chip assay (Agilent Technologies, Santa Clara, CA, USA), respectively. The libraries were sequenced using PacBio RSII and Sequel instruments. On the PacBio RSII instrument, libraries were sequenced with SMRT bell-Polymerase Complex using the P6 DNA/Polymerase binding kit 2.0 (Pacific Biosciences, Menlo Park, CA, USA) according to the manufacturer's instructions. The complex was loaded into a SMRT cell v3.0 (Pacific Biosciences), recording a series of microscope pictures for a period of six hours. On the PacBio Sequel instrument, the libraries were sequenced with SMRT bell-Polymerase Complex created using the Sequel binding kit 1.0 (Pacific Biosciences) according to the manufacturer instructions. The PacBio Sequel instrument was programmed to load the sample on a Sequel SMRT Cell 1M (Pacific Biosciences). The SMRT cell was sequenced recording a series of microscope pictures for a period of ten hours with a Sequel Sequencing 1.21 chemistry (Pacific Biosciences). For both the RSII and Sequel, a MagBead loading V2 (Pacific Biosciences) method was used to enrich for longer fragments. The complete PacBio sequence dataset was *de novo* assembled with FALCON Unzip, v.0.4.0 (Chin, et al. 2016) using a plant-specific configuration file (fc_run_plant.cfg). For polishing and genome size estimation, the same gDNA sample of 'Gala Galaxy' used for PacBio library preparation was also used to prepare Illumina libraries (Illumina TruSeq Nano DNA Library Preparation, Illumina, San Francisco, CA, USA) with an average insert size of 500 bp. Arrow was the polishing option chosen when applying the variantCaller v.2.3.3 implementation bundled in SMRT Link v 5.1.0. The genome size was estimated using Jellyfish 2.2.6 (Marçais and Kingsford 2011) and the estimate_genome_size script (https://github.com/josephryan/estimate_genome_size.pl) and the forward reads (R1) of the Illumina sequenced library of ‘Gala Galaxy’.

**Hybrid scaffolding**

To increase the contiguity and to phase the contigs generated by FALCON Unzip, two optical maps were constructed using different labelling kits. These were combined to generate dual enzyme hybrid scaffolds. To generate the optical maps, DNA was extracted from four grams of apple leaves of ‘Gala Galaxy’. These samples were prepared with the Plant DNA Isolation kit (Bionano Genomics, San Diego, CA, USA) following the Bionano Prep Plant Tissue DNA Isolation Base Protocol (Document Number 30068 Rev C). The Bionano Irys and Saphyr platforms generated optical maps using DNA labeled with DLS and NLRS kits, respectively. For DLS, DNA was labeled using the Bionano Prep DNA Labeling Kit-DLS (Bionano Genomics) according to manufacturer’s instructions. A total of 750 ng of purified genomic DNA was labeled with DLE labeling Mix and subsequently incubated with Proteinase K (Qiagen, Hilden, DE) followed by drop dialysis. After the clean-up step, the DNA was pre-stained, homogenized, and quantified using on a Qubit Fluorometer to establish the appropriate amount of backbone stain. The staining reaction was incubated at room temperature for at least two hours. For NLRS, DNA was labeled according to manufacturer's instructions using the Prep DNA Labeling Kit-NLRS (Bionano Genomics,). Three-hundred nanograms of purified genomic DNA were treated with Nb.BspQI (New England Biolabs, Ipswich, MA, USA) in NEB Buffer 3. The nicked DNA was labeled with a fluorescent-dUTP nucleotide analog using Taq DNA polymerase (New England BioLabs). After labeling, nicks were repaired with Taq DNA ligase (New England BioLabs) in the presence of dNTPs. The backbone of fluorescently labeled DNA was counterstained overnight with YOYO-1 (Bionano Genomics). The *de novo* assembly of the optical maps was performed using the Bionano Access v1.2.1 and Bionano Solve v3.2.1 software. The assembly type performed was the "haplotype" with "no extend split" and "no cut segdups". Default parameters were adjusted to accommodate the genomic properties of the apple genome. Specifically, the minimal length for the molecules to be used in the assembly was 150 kb, the "Initial P-value" cut off threshold was adjusted to $1 \times 10^{-11}$ and the P-value cut off threshold for extension and refinement was set to $1 \times 10^{-12}$ according to manufacturer's guidelines.

**Scaffolds anchoring to a genetic map and to a previously reported haploid genome**

Sequences flanking the SNP markers from the genetic map of 'Fuji' × 'Gala' (Di Pierro, et al. 2016) were BLAST searched against the hybrid assembly, retaining only the hits with highest bit score. Physical position and genetic position of each SNP served as input for ALLMAPS anchoring. In addition, the assembly was anchored to the doubled-haploid reference GDDH13 assembly (Daccord, et al. 2017) by means of aligning GDDH13's coding sequences through BLAST

**RNA extraction and library preparation.**

RNA was extracted using the Quick-RNA MiniPrep Kit (Zymo Research, Irvine, CA, USA) following the manufacturer's protocol with the exception that RNA was eluted twice with 35µl DNAse free water. Residual DNA was digested with 1 µl DNAseI (DNA-free DNA removal kit, Thermo Fisher Scientific, Waltham, MA, USA). Complementary RNA libraries were created using the TrueSeq®RNA (Illumina, San Diego, CA, USA) kit, including a polyA purification step according to manufacturer protocol.

**Genome visualization for coverage and phase analysis of the assembled chromosomes**

To further evaluate the correctness of the genome assembly, diagnostic genome features were visualized in RStudio (Version 1.2.5001) using the karyoploteR package (Gel and Serra 2017). Illumina read coverage, genetic phase, and correlation of genetic and physical order on the assembly were assessed within and across pseudomolecules. Furthermore, homologous regions between each haploid assembly were visualized. By inspecting the Illumina read coverage distribution, homozygous regions in the genome, in which both haplotypes were collapsed to a single sequence, were identified. Such regions displayed a coverage two-fold higher when compared to regions assembled in two separate haplotypes. The coverage of the assembly by Illumina reads was calculated from the bam-file generated by mapping the Illumina paired-end reads using the “Map to reference function” in CLC genomics workbench 11.0 (Length fraction = 0.8, Similarity fraction = 0.95 and ignoring non-specific matches). Phase information published with the genetic map of 'Fuji' × 'Gala' (Di Pierro, et al. 2016) was used to determine the phase of each SNP in the two haploid assemblies (therefore using only SNP markers heterozygous in 'Gala'). The two sequences corresponding to the two alleles in ‘Gala’ for each SNP locus were generated and BLAST-searched separately on the primary and secondary haploid assemblies. BLAST hits were then parsed to allocate the polymorphisms to the primary and secondary haploid assemblies. The SNPs BLAST hits used for ALLMAPS-based anchoring served also for generating the links between primary and secondary assembly. In addition, the genetic position (in cM) of the SNP markers was included in order to visualize the collinearity of genetic and physical map.

**Graph-based assembly**

The primary haploid assembly MDGGph_v1.0 was used as haploid reference for graph-based (WhatsHap) assembly of the genome (Patterson, et al. 2015) with default settings. Pacbio and Illumina reads were mapped to the reference genome and WhatsHap generated two versions of the haploid reference that should correspond to the two phased haploid genomes. The KAT was used as described above to evaluate the completeness of the resulting diploid phased genome in comparison with the GDDH13 assembly and the unphased diploid assembly MDGGdi_v1.0
