## Supplementary Figures and Tables for "Chromosome-scale *de novo* diploid assembly of the apple cultivar ‘Gala Galaxy’"

Supplementary figure 1: Syntenic dotplot generated comparing the 17 apple chromosomes (chr1-chr17) of the GDDH13 assembly (x-axis, Daccord et al 2017) versus the primary haploid assembly MDGGph_v1.0 (y-axis). Each dot indicates a syntenic gene pair between the two assemblies.

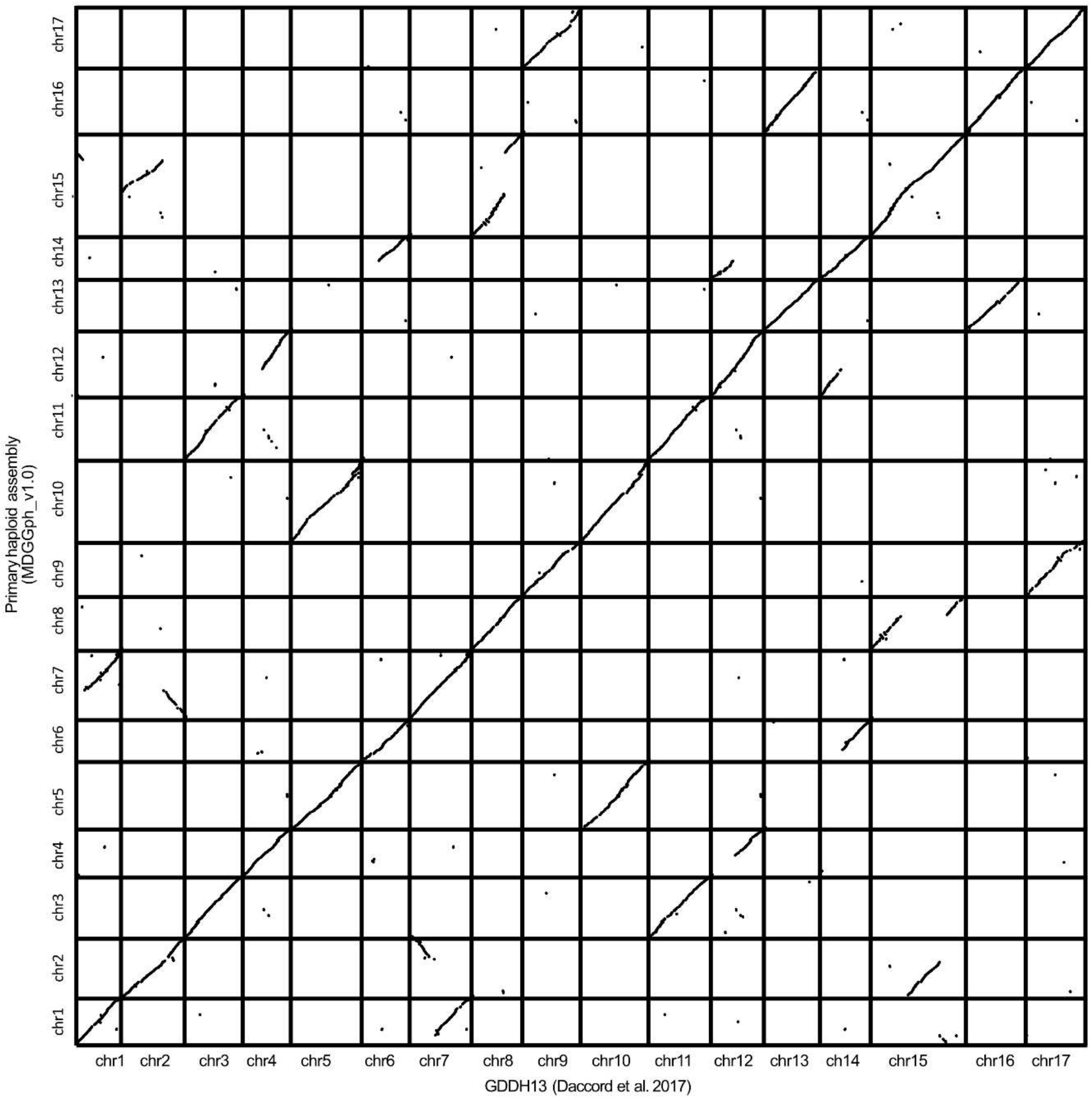

Supplementary table 1: Chromosome lengths of the haploid assemblies MDGGph_v1.0 and MDGGsh_v1.0 compared to HFTH1 (Zhang et al. 2019) and GDDH13 (Daccord et al 2017).

| **MDGGph_v1.0** (primary haploid assembly) | | | **MDGGsh_v1.0** (secondary haploid assembly) | | | **HFTH1** (Zhang et al. 2019) | | | **GDDH13** (Daccord et al 2017) | | |
|---|---|---|---|---|---|---|---|---|---|---|---|
| | **Length (bp)** | **Scaffolds** | | **Length (bp)** | **Scaffolds** | | **Length (bp)** | **Scaffolds** | | **Length (bp)** | **Scaffolds** |
| **chr1** | 34,957,745 | 2 | chr_s1 | 20,900,120 | 189 | chr1 | 32,944,118 | 13 | chr1 | 32,625,452 | 86 |
| **chr2** | 43,219,086 | 2 | chr_s2 | 42,574,754 | 180 | chr2 | 38,449,405 | 9 | chr2 | 37,577,729 | 88 |
| **chr3** | 41,865,124 | 2 | chr_s3 | 35,827,395 | 240 | chr3 | 37,138,690 | 5 | chr3 | 37,524,076 | 81 |
| **chr4** | 36,846,674 | 2 | chr_s4 | 26,433,372 | 183 | chr4 | 31,012,745 | 8 | chr4 | 32,301,874 | 65 |
| **chr5** | 46,262,512 | 3 | chr_s5 | 58,045,489 | 239 | chr5 | 47,891,858 | 14 | chr5 | 47,952,461 | 108 |
| **chr6** | 37,227,775 | 3 | chr_s6 | 46,706,699 | 155 | chr6 | 35,567,198 | 6 | chr6 | 37,137,259 | 89 |
| **chr7** | 42,138,739 | 3 | chr_s7 | 23,675,059 | 246 | chr7 | 35,934,761 | 6 | chr7 | 36,691,129 | 76 |
| **chr8** | 37,261,338 | 1 | chr_s8 | 25,480,337 | 200 | chr8 | 31,511,015 | 8 | chr8 | 31,609,270 | 67 |
| **chr9** | 42,757,514 | 1 | chr_s9 | 30,211,764 | 217 | chr9 | 34,800,404 | 10 | chr9 | 37,604,908 | 80 |
| **chr10** | 49,447,705 | 2 | chr_s10 | 46,315,672 | 241 | chr10 | 43,815,736 | 14 | chr10 | 41,762,413 | 83 |
| **chr11** | 46,034,967 | 3 | chr_s11 | 48,391,545 | 218 | chr11 | 42,456,296 | 15 | chr11 | 43,059,885 | 91 |
| **chr12** | 37,592,879 | 1 | chr_s12 | 19,120,998 | 191 | chr12 | 32,285,079 | 9 | chr12 | 33,050,054 | 75 |
| **chr13** | 40,714,413 | 1 | chr_s13 | 37,617,649 | 237 | chr13 | 44,866,511 | 13 | chr13 | 44,339,518 | 119 |
| **chr14** | 36,122,606 | 1 | chr_s14 | 39,177,835 | 148 | chr14 | 31,515,206 | 6 | chr14 | 32,513,452 | 62 |
| **chr15** | 60,319,455 | 2 | chr_s15 | 48,141,362 | 339 | chr15 | 56,644,392 | 16 | chr15 | 54,945,402 | 129 |
| **chr16** | 47,681,219 | 1 | chr_s16 | 28,833,803 | 228 | chr16 | 41,670,059 | 15 | chr16 | 41,389,449 | 93 |
| **chr17** | 39,000,747 | 1 | chr_s17 | 23,971,657 | 211 | chr17 | 33,998,825 | 8 | chr17 | 34,748,701 | 76 |
| | | | Unanchored | 80,621,795 | 2583 | Unanchored | 7,992,922 | | Unanchored | 52,728,359 | |

Supplementary table 2: Results of the Benchmarking Universal Single-Copy Orthologs (BUSCO) analysis on the assemblies MDGGph_v1.0, MDGGdi_v1.0 and on GDDH13 (Daccord *et al.* 2017).

| | **MDGGph_v1.0 (primary haploid assembly)** | | **MDGGdi_v1.0 (diploid assembly)** | | **GDDH13 (Daccord et al. 2017)** | |
|---|---|---|---|---|---|---|
| **Complete BUSCOs (C)** | 1,347 | 93.5% | 1,387 | 96.3% | 1,383 | 96.0% |
| **Complete and single-copy BUSCOs (S)** | 943 | 65.5% | 467 | 32.4% | 894 | 62.1% |
| **Complete and duplicated BUSCOs (D)** | 404 | 28.1% | 920 | 63.9% | 489 | 34.0% |
| **Fragmented BUSCOs (F)** | 20 | 1.4% | 11 | 0.8% | 13 | 0.9% |
| **Missing BUSCOs (M)** | 73 | 5.1% | 42 | 2.9% | 44 | 3.1% |
| **Total BUSCO groups searched** | 1,440 | | 1,440 | | 1,440 | |
